# Uncovering phenotypic diversity and cell-type-specific prefrontal cortex calcium dynamics in rat fentanyl self-administration

**DOI:** 10.64898/2026.05.28.728535

**Authors:** Alexander C. Whitebirch, Shawn Panh, Loveleen Tripathi, Aaron F. Garcia, Nailyam Nasirova, Dayna Suess, Susan M. Ferguson

## Abstract

**BACKGROUND:** The proliferation of the potent synthetic opioid fentanyl has exacerbated the ongoing crisis of substance use disorder and associated overdose deaths, yet the neurobiological mechanisms that underlie individual vulnerability to addiction and relapse remain poorly understood, particularly in the context of fentanyl use. The prefrontal cortex (PFC) has been identified as a key brain structure important for cognitive functions impacted in addiction, including inhibitory control of behavior and association of drug experience with specific cues, contexts, or actions. Although the heterogenous neuronal composition of the PFC complicates attribution of addiction-related behavioral regulation to specific cortical cell types and circuits, application of cell-type-specific methods in translationally relevant rodent models have begun to elucidate the key neural substrates of opioid addiction.

**METHODS:** We used an intermittent access fentanyl self-administration (IntA SA) model to characterize individual variation and sex differences in addiction vulnerability in male and female rats. Longitudinal wireless fiber photometry recording was used to track calcium activity patterns in intratelencephalic (IT) neurons of the prelimbic cortex across acquisition of self-administration, escalation of fentanyl intake, extinction training, and cue-induced reinstatement of fentanyl seeking.

**RESULTS:** We found that our fentanyl IntA SA paradigm produces distinct low- and high-risk addiction severity phenotypes and that female rats exhibited a greater propensity for high-risk classification, which was characterized by abundant and consistent fentanyl intake, robust responsiveness to conditioned and discriminative fentanyl-associated cues, and high levels of fentanyl-seeking during periods of drug unavailability, extinction training, and a cue-induced reinstatement test. Fiber photometry recordings revealed dynamic encoding of fentanyl-associated stimuli by prelimbic IT neurons across the IntA SA paradigm with event-related calcium transients observed in association with lever presses, fentanyl infusions, and presentation of conditioned and discriminative cues.

**CONCLUSIONS:** Our data indicate that fentanyl IntA SA is a translationally relevant paradigm that enables investigation of phenotypic diversity and the role of sex in fentanyl addiction. Longitudinal cell-type-selective calcium recordings revealed dynamic representation of fentanyl-associated stimuli by IT neurons of the prelimbic cortex consistent with a role for this cortical subpopulation in addiction-related behaviors.

## INTRODUCTION

Recent increases in opioid use disorder (OUD) prevalence have been driven, in part, by the proliferation of the potent synthetic opioid fentanyl^1–5^. Although fentanyl users are at high risk for relapse and overdose^1,6–11^, few studies have examined the neurobiological consequences of chronic fentanyl use. Given fentanyl’s profound impact on the landscape of illicit drug use and overdose, understanding the specific brain structures and circuits that mediate fentanyl addiction is critical. Notably, only a subset of individuals who use drugs develop substance use disorders^12^, and the genetic and neurobiological mechanisms underlying this variability remain incompletely understood^13^. Among opioid users, women report more severe withdrawal^14^ and greater cue-induced craving^15^. Accordingly, it is vital that preclinical investigations examine phenotypic diversity and sex differences in addiction-like behaviors using translationally-relevant animal models^16–20^.

Studies in individuals with chronic drug use^21–25^ and in animal models^26–32^ have consistently implicated the prefrontal cortex (PFC) in addiction. Common findings include reduced basal activity of PFC neurons along with enhanced PFC responsiveness to drug self-administration and drug-associated cues, reflecting the established role for PFC in inhibitory control of behavior and cue-reward processing. Within the rodent PFC, the prelimbic region regulates drug-seeking behaviors by encoding associations between cues, contexts, or actions and subsequent drug intake, thereby supporting information processing related to operant contingencies or punishment^33,34,43,44,35–42^.

Identifying the role of discrete cortical cell populations and circuits in addiction-related behaviors is complicated by the heterogeneity of cortical networks. The prelimbic cortex contains diverse sets of projection neurons with distinct anatomical and physiological characteristics^45–47^, yet few studies have leveraged cell-type-selective approaches to resolve the roles of these subpopulations, with most relying on non-selective pharmacological manipulations^48–53^. Intratelencephalic (IT) neurons receive widespread synaptic input from many brain regions, including the claustrum, thalamus, globus pallidus, hypothalamus, and ventral pallidum, and in turn establish dense bilateral projections predominately to the cortex and striatum (i.e., dorsal striatum and nucleus accumbens, NAc). In contrast, pyramidal tract (PT) neurons receive limited input primarily from the cortex but project widely throughout the brain with extensive collaterals that reach subcortical targets including the thalamus, subthalamic nucleus, ventral tegmental area, rostromedial tegmental nucleus, and locus coeruleus^47,54^. Although some PT neurons provide input to ipsilateral striatum alongside IT neurons, retrograde tracing studies indicate that the cortical projections to NAcC and to extratelencephalic targets such as VTA or RMTg arise from distinct, largely nonoverlapping neuronal subpopulations, potentially reflecting further heterogeneity within the PT population^55,56^. IT neurons provide input to PT neurons, yet receive relatively sparse PT input in return^45,57^. Thus, while extensive evidence supports a critical role for the corticostriatal pathway in addiction-related behaviors, particularly cue-induced reinstatement of drug-seeking^40,48,62,50–53,58–61^, the respective contributions of IT and PT neurons remain unknown. Taken together, these findings suggest that IT neuronal ensembles are poised to receive and integrate drug-associated information from a wide array of sources and in turn orchestrate drug-taking and drug-seeking through recruitment of intracortical and striatal targets.

To examine individual- and sex-specific variability in fentanyl addiction, we used an intermittent access self-administration model (IntA SA), which we have previously shown generates distinct low- and high-risk addiction-like phenotypes with heroin across multiple behavioral metrics^63,64^. In a subset of animals, we examined the role of the IT subpopulation of prelimbic neurons in fentanyl self-administration using an intersectional viral strategy that enables cell-type-specific monitoring of neural activity in rats. Fiber photometry recordings of calcium signals from prelimbic IT neurons were performed across acquisition, escalation, extinction, and cued reinstatement of fentanyl-seeking and neuronal activity was correlated with behavioral performance.

## METHODS AND MATERIALS

### Animal use

Outbred Sprague-Dawley rats weighing 175 - 275 g upon arrival (n = 27 males, 24 females, Envigo) were dual-housed in a temperature- and humidity-controlled, specific-pathogen free colony room and were acclimated on a 12:12 h light:dark cycle with *ad libitum* food/water. Behavioral procedures typically began in the morning shortly after the beginning of the dark cycle. The University of Washington’s IACUC provided oversight and approval of all procedures.

### Drugs

Fentanyl HCl was obtained through the Drug Supply Program of the National Institute on Drug Abuse (NIDA) and dissolved in sterile saline (0.9% NaCl) to a stock concentration of 0.25 mg/mL. Syringes were prepared based on body weight for administration of 0.0015 mg/kg per infusion, with a volume of 50 μL delivered over 2.8 seconds.

### Viral vectors

A Cre-dependent GCaMP6s viral vector (AAV9-CAG-FLEX-GCaMP6s-WPRE-SV40) was obtained from Addgene (100842-AAV9) at a titer of 1.9x10^13^ viral genomes (vg) per mL. An AAV8-CaMKIIa-eGFP vector was prepared by Dr. Selena Schattauer in the University of Washington Center for the Neurobiology of Addiction, Pain, and Emotion (NAPE) Molecular Genetics Research Core (MGRC) at a titer of 4.9x10^12^ (commercially available as Addgene 50469-AAV8). Retrograde recombinase delivery was accomplished using AAVrg-EF1α-Cre obtained from Addgene (55636-AAVrg) at a titer of 2.5x10^13^.

### Retrograde tracing with cholera toxin subunit B

Retrograde tracing to selectively label prefrontal IT or PT neurons was accomplished using cholera toxin subunit B (CTB) conjugated to Alexa Fluor 594 (Thermo Fisher C-34777) or 647 (Thermo Fisher C-34778). 500 nL of CTB solution (0.5 – 1% w/v in sterile saline) was infused using stereotaxic procedures (see below) into contralateral NAcC (for IT neurons) or ipsilateral pyramidal tract in the pons (for PT neurons). Twenty-eight days following infusion animals were perfused transcardially perfused with phosphate buffered saline (PBS) followed by 4% paraformaldehyde in PBS and brains prepared for imaging as described below.

### Surgical procedures

Rats were anesthetized with isoflurane (1 – 5% inhaled; Patterson Veterinary) and provided with meloxicam (1 mg/mL, 1 mg/kg subcutaneous injection; Patterson Veterinary) for analgesia. Heat support was provided using a warmed water blanket (HTP-1500). All surgical sites were scrubbed with alternating povidone-iodine and ethanol, and eye lubrication was applied at the beginning of surgery. Rats were provided with subcutaneous saline (0.9% NaCl) and monitored for at least 3 days following each surgical procedure, and recovered for at least a week between procedures.

#### Stereotaxic surgeries

Stereotaxic surgeries were performed following standard procedures and a Kopf stereotaxis frame and a digital stereotaxic coordinate system (Kopf Model 942). Viral vectors and CTB solutions were infused using gas-tight syringes (10 µL, 7635-01, Hamilton Company) with 33-gauge needles driven by motorized injectors (World Precision Instruments UMP3) and controller units (World Precision Instruments Micro2T). Craniotomies were drilled using a handheld micromotor (Foredom MH-130). Stereotaxic coordinates were measured relative to the skull surface at bregma and were determined through review of the literature, reference to the Paxinos and Watson rat brain atlas^65^, and adjusted as needed following pilot experiments: prelimbic cortex, A-P +2.8, M-L -0.64, D-V -4.2:-3.8; contralateral nucleus accumbens core (NAcC), A-P +1.5, M-L 1.38, D-V -7.2:-6.6; pons, A-P -9.2, M-L -2.33, D-V -10.55:-10.45 (10-degree lateral angle). For photometry experiments, 2000 nL of AAVrg-EF1α-Cre was infused into NAcC_contra_ for a dose of 5x10^9^ vg, and 2000 nL of either AAV9-CAG-FLEX-GCaMP6s-WPRE-SV40 (dose range of 5.9x10^8^ to 2.4x10^9^) was infused into the prelimbic cortex. Following virus infusion, fiber optic cannula (MFC_400/430-0.66_5.5mm_ZF2.5(G)_FLT, Doric) were lowered into place in the prelimbic cortex (0.1 mm above the center of virus infusion) using a holder affixed to the stereotax arm. Needles were left in place for 10 minutes after virus or CTB infusion to allow for diffusion away from the injection site.

#### Catheter implant surgeries

Jugular catheterization surgeries were performed as previously described (O’Neal et al., 2020). Chronic indwelling catheters were placed into the right jugular vein and attached to a back-mounted port. Catheters were flushed daily with a solution of gentamicin sulfate (5 mg/mL, Patterson Veterinary) and heparin (3 U/mL, Patterson Veterinary) to maintain patency and minimize infection. Catheter patency was tested before self-administration and then regularly throughout the experiment through infusion of brevital sodium (10 mg/mL in sterile saline, 0.05–0.2 mL/ infusion, iv; methohexital sodium, Patterson Veterinary). Catheter patency was confirmed if rats became ataxic within 5 seconds of infusion. Rats that lost patency prior to the beginning of extinction training (n= 4 males, 5 females) were excluded from all analysis.

### Fentanyl self-administration

Rats were trained to self-administer fentanyl in 60-minute sessions during which presses on an active lever were rewarded on a fixed ratio of 1, with each lever press resulting in a 4 second cue light illumination and infusion of fentanyl (0.0015 mg/kg over 2.8 seconds, maximum of 10 infusions). Presses on an inactive lever were recorded but produced no reward or cue. Rats were trained for 5-7 days with no significant change in performance beyond 5 days. Rats quickly learned to discriminate between the active and inactive lever and most animals earned the maximum number of daily infusions within 5 days of training (9.34 ± 0.25 infusions on day 5; Supplemental Figure 1). Rats then underwent 10 days of intermittent access fentanyl self-administration (IntA SA). Each day of IntA SA consisted of 335 mins of alternating drug available (DA, 5 min) and drug unavailable (DU, 25 min) periods. Continuous illumination of the houselight was used as a discriminative cue signaling fentanyl unavailability, and levers remained available throughout the experiment to monitor fentanyl-seeking. During unavailable periods, active lever presses were unrewarded (i.e., no cue light illumination or fentanyl infusion). Houselights were extinguished at the onset of each available period, during which presses on the active lever yielded cue light illumination and fentanyl infusion. A progressive ratio (PR) test was performed to assess motivation for fentanyl under an increasing schedule of reinforcement (1, 2, 4, 6, 9, 12, 15, 20, 25, 32, 40, 50, 62, 77, 95, 118, 145, 178, 219, 268, 328, 402, 492, 603). The PR test had a maximum duration of 6 hours and concluded when 60 min passed without an earned infusion. Following the PR test, rats underwent three additional days of IntA SA to re-baseline behavior. The extinction phase of the experiment consisted of daily 1 hour sessions during which presses on both the active and inactive levers yielded neither cues nor fentanyl infusions. A cued reinstatement (CR) test was performed following extinction training. The cue light was illuminated for 10 seconds at the onset of this 60 min test, after which active lever presses resulted in illumination of the cue light (but no fentanyl infusion). In photometry experiments, rats then underwent 6 days of re-extinction followed by a second CR test. For these animals, only the first 10 days of extinction and first CR test were included in behavioral analyses.

### Addiction severity classification

Eight measures were used to quantify the severity of addiction-like behavior: (1) *acquisition*, the number of days required for an animal to earn 10 infusions; (2) *infusion latency*, the average duration in seconds before the first infusion of each available period; (3) *intake*, the total number of fentanyl infusions earned over the first 10 days of IntA SA; (4) *consistency*, the percentage of available periods over the first 10 days of IntA SA in which at least one infusion was earned; (5) *seeking*, the total number of active lever presses occurring during unavailable periods over the first 10 days of IntA SA; (6) *motivation,* the total number of active lever presses during the PR test; (7) *extinction*, the total number of active lever presses occurring over 10 days of extinction training; (8) *reinstatement*, the total number of active lever presses occurring during the cued reinstatement test. The raw values for each measure were converted to z-scores (subtraction of the group mean and division by the standard deviation), and individual z-scores were summed within each animal into cumulative z-scores used to classify rats as low-risk (sum < -1), moderate (sum between -1 and +1), or high-risk (sum > +1).

### Fiber photometry

Fiber photometry recordings were performed using a wireless photometry system (Doric), in which wireless headstages (WiFP_IE415_E470_F520_2.5mm_A) with integrated photodetectors and antenna transmitted real-time fluorescence measurements to a base station that interfaced with Doric Neuroscience Studio (version 6.6.3.9) software via the Doric Neuroscience Console 500. GCaMP6s fluorescence was detected using a 470 nm wavelength, and a 415 nm wavelength was used as an isosbestic control for all recordings. LED power was set to deliver a light intensity of approximately 40 μW. Recordings were performed at all phases of the self-administration paradigm: training, IntA SA, PR test, extinction, and CR test. During the training, extinction, and CR phases, recordings were continuous throughout the sessions. To avoid excessive photobleaching during the IntA SA phase, in which daily sessions lasted 335 mins, we recorded multiple 15-min intervals that encompassed drug available periods along with flanking late and early drug unavailable periods.

### Immunohistochemistry and imaging

Rats were euthanized with Beuthanasia-D (pentobarbital sodium/phenytoin sodium, Patterson Veterinary) and transcardially perfused with chilled phosphate-buffered saline (PBS) followed by 4% paraformaldehyde (PFA). Brains were removed, immersed in 4% PFA overnight at 4 degrees C, and stored in PBS thereafter. For rats implanted with optic fibers, heads were left intact with headcaps in place for the overnight post-fixation to ensure the fiber tracts were firmly fixed, and the implants were removed and brains extracted the following day. Brains were sectioned at 40 μm using a Leica VT1200 vibrating blade microtome and sections were stored in PBS at 4 degrees C. To examine endogenous fluorescence from viral expression or Alexa Fluor-conjugated CTB, sections were then mounted onto slides, allowed to air dry, and then coverslipped with DAPI mounting medium (Vector Laboratories). To amplify endogenous fluorescence from GCaMP6s, sections were first permeabilized in 0.5% Triton X-100 in PBS for 10 minutes at room temperature on a shaker. Sections were then placed in a blocking solution of 0.25% Triton X-100 and 5% normal goat serum (NGS) in PBS for 2 hours at room temperature on a shaker. Blocking solution was then removed and a primary antibody solution was applied, which consisted of 0.25% Triton X-100, 2.5% NGS, and mouse anti-GFP (1:400, Takara Living Colors 632375) in PBS. Sections were incubated in the primary antibody solution overnight on a shaker at 4 degrees C. Sections were washed 3 times for 10 minutes/wash in 0.25% Triton X-100 in PBS, and then were incubated in a secondary antibody solution of 2.5% NGS and either goat anti-mouse conjugated to Alexa Fluor 488 (1:400, Invitrogen A11001) for 4 hours on a shaker at room temperature. Finally, sections were washed 3 times for 10 minutes/wash in 0.25% Triton X-100 in PBS, stored in PBS, and mounted onto slides as previously described. Images were captured using a Zeiss apotome microscope (Imager.M2) and Zen Blue software (version 2.6) and processed using ImageJ/FIJI (version: 2.16.0/1.54p).

### Data analysis and statistics

Behavioral data were collected using Med Associates software (MED-PC V version 5.1), processed with custom-written Python scripts (version 3.11.3), and statistical analyses were performed using GraphPad Prism 10 (version: 10.6.1). Comparisons incorporating multiple groups and timepoints were performed using two- or three-way repeated-measures analysis of variance (RM ANOVA) with the Geisser-Greenhouse correction and Šídák’s multiple comparisons test, or, where appropriate, a Fisher’s least significant difference test with manual Bonferroni correction. In cases where datasets had missing values, Prism applies a comparable mixed-effects model, which utilizes a compound symmetry covariance matrix fit using Restricted Maximum Likelihood (REML). Cross-group comparisons (e.g. of the severity classification metrics in Figure 1) were performed using Welch’s t tests or Kruskal-Wallis tests, while within-group tests (e.g. of lever press totals at early and late timepoints in Figure 3) were performed using paired t tests. Comparison of data median against a hypothetical mean of 0 was performed using the Wilcoxon signed-rank test. Comparisons of lever press / infusion escalation or decline across time were performed using simple linear regression. For all comparisons, α ≤ 0.05. Pre-processing and analysis of photometry data was performed using a custom-written pipeline in Python informed by published protocols^66,67^. Pre-processing consisted of several successive steps: signal downsampling to 20 Hz and low-pass filtering with a cutoff of 8 Hz; artifact detection and exclusion using linear interpolation; alignment of photometry signals to behavioral event timestamps; motion correction by linear regression of the GCaMP and isosbestic channels; photobleaching and baseline drift correction using subtraction of a double exponential fit and a baseline computed with an adaptive iteratively reweighted penalized least squares (airPLS) algorithm^68^; z-score normalization of the GCaMP and isosbestic signals. Perievent fluorescence was aligned to active and inactive lever presses, fentanyl infusions, missed infusions in extinction and CR sessions, period transitions in IntA SA sessions, and lever extension at session initiation. Normalized fluorescence amplitudes and area under the curve values (computed via trapezoidal numerical integration) were measured within specified time windows for both spontaneous calcium transients and perievent signals. Post hoc statistical analysis of peri-event calcium activity was performed using the Fiber Photometry Post Hoc Analysis (FiPhoPHA) Python package^69^. Event-related transients (ERTs) were identified using a bootstrapped confidence interval (bCI), a resampling method that provides temporally precise, nonparametric, and unbiased identification of intervals during which a photometry signal significantly deviates from the null hypothesis, a baseline value of 0, or from a comparison signal^70^. This method effectively reduces Type I and Type II error rates while providing statistically powerful and relatively unbiased analysis of peri-event signals in fiber photometry data^70,71^. Comparisons across timepoints or groups were also assessed using non-parametric resampling-based permutation tests in the FiPhoPHA package. We used a bCI threshold of 95%, 2000 resamples and permutations, and a consecutive threshold of 1000 ms to identify ERTs and to test for differences across timepoints or groups.

**Figure 1.**
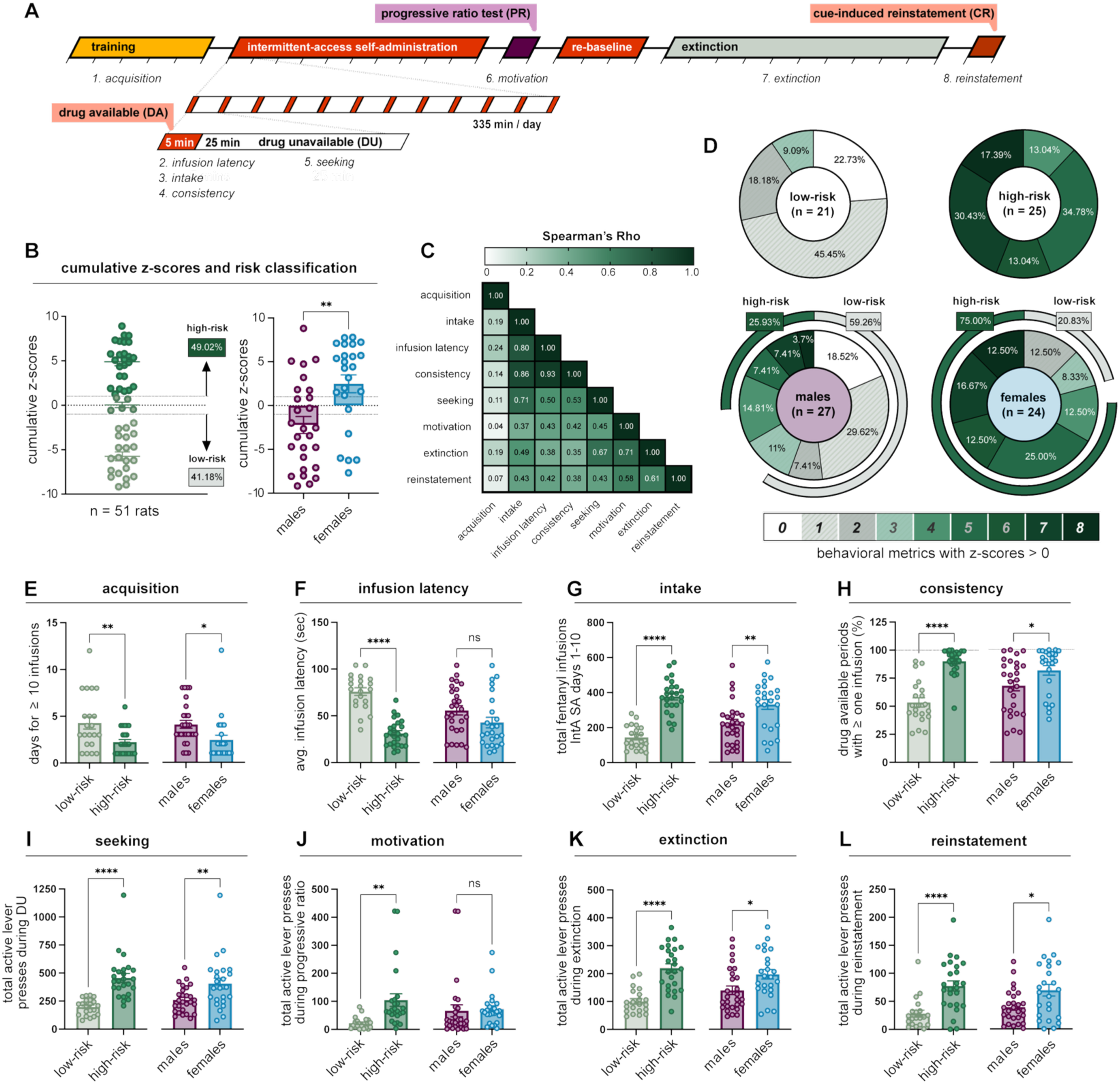
Intermittent access fentanyl self-administration (IntA SA) reveals low-risk and high-risk behavioral phenotypes and sex differences in addiction severity. **(A)** Timeline of the fentanyl IntA SA self-administration paradigm. **(B)** Cumulative z-scores reflecting the severity of addiction-like behaviors were used to classify rats as either low-risk (41.18% of animals) or high-risk (49.02% of animals). Female rats exhibited higher cumulative z-scores than males. **(C)** Behavioral metrics used for addiction severity classification were highly correlated. **(D)** High-risk rats scored positive on more severity metrics as compared to low-risk rats. Similarly, female rats scored positive on more metrics than males, and a greater proportion of females were classified as high-risk. **(E, G – I, K, L)** Relative to low-risk and male rates, high-risk and female rats reached an intake criterion of 10 daily fentanyl infusions in fewer days **(E)**, earned a greater total number of fentanyl infusions **(G)**, earned fentanyl infusions more consistently during IntA SA sessions **(H)**, and exhibited higher levels of active lever pressing during periods of fentanyl unavailability **(I)**, extinction sessions **(K)**, and the cued reinstatement test **(L)**. **(F)** During available periods in the IntA SA sessions, high-risk rats earned infusions with a shorter latency as compared to low-risk animals. There was no significant difference in infusion latency between males and females. **(J)** High-risk rats pressed the active lever a greater number of times during the PR test as compared to low-risk rats. There was no significant sex difference in PR lever pressing. N = 51 rats total, 21 low-risk, 25 high-risk, 27 male, 24 female.

**Figure 2.**
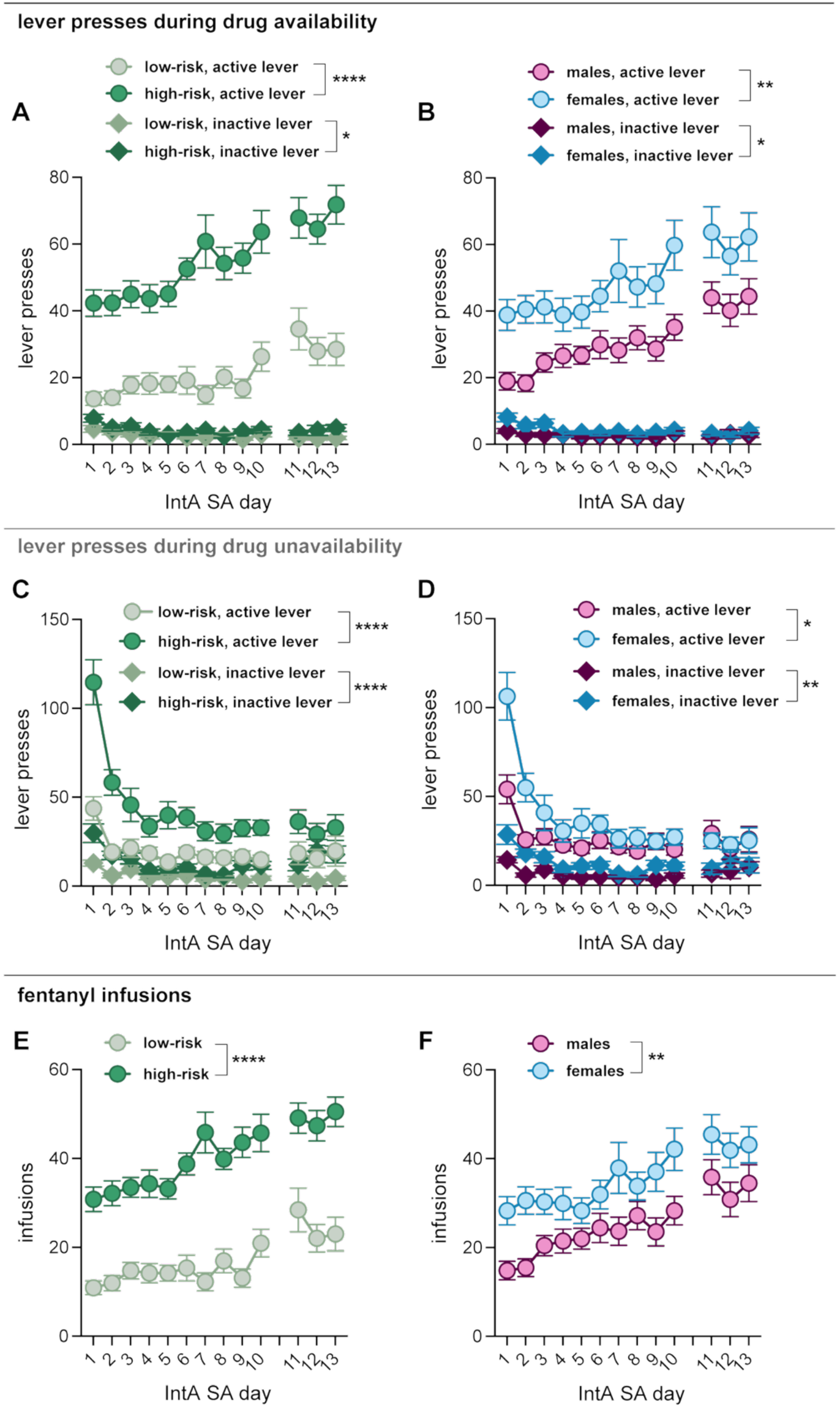
Escalation of fentanyl self-administration across sexes and addiction severity phenotypes. Over the course of 13 days of fentanyl IntA SA, rats gradually increased their daily active lever pressing and displayed steady minimal presses of the inactive lever. **(A, B)** High-risk and female rats had higher daily active and inactive lever presses than low-risk rats and female rats, respectively. **(C, D**) Over the course of IntA SA, lever pressing during unavailable periods rapidly decreased, following a substantial number of active lever presses that occurred on the first IntA SA day. High-risk and female rats had a higher level of both active lever and inactive lever pressing during unavailable periods as compared to low-risk and female rats, respectively. **(E, F)** Rats gradually escalated their fentanyl intake over the course of 13 days of IntA SA. High-risk and female rats had greater levels of daily intake as compared to low-risk and male rats, respectively.

**Figure 3.**
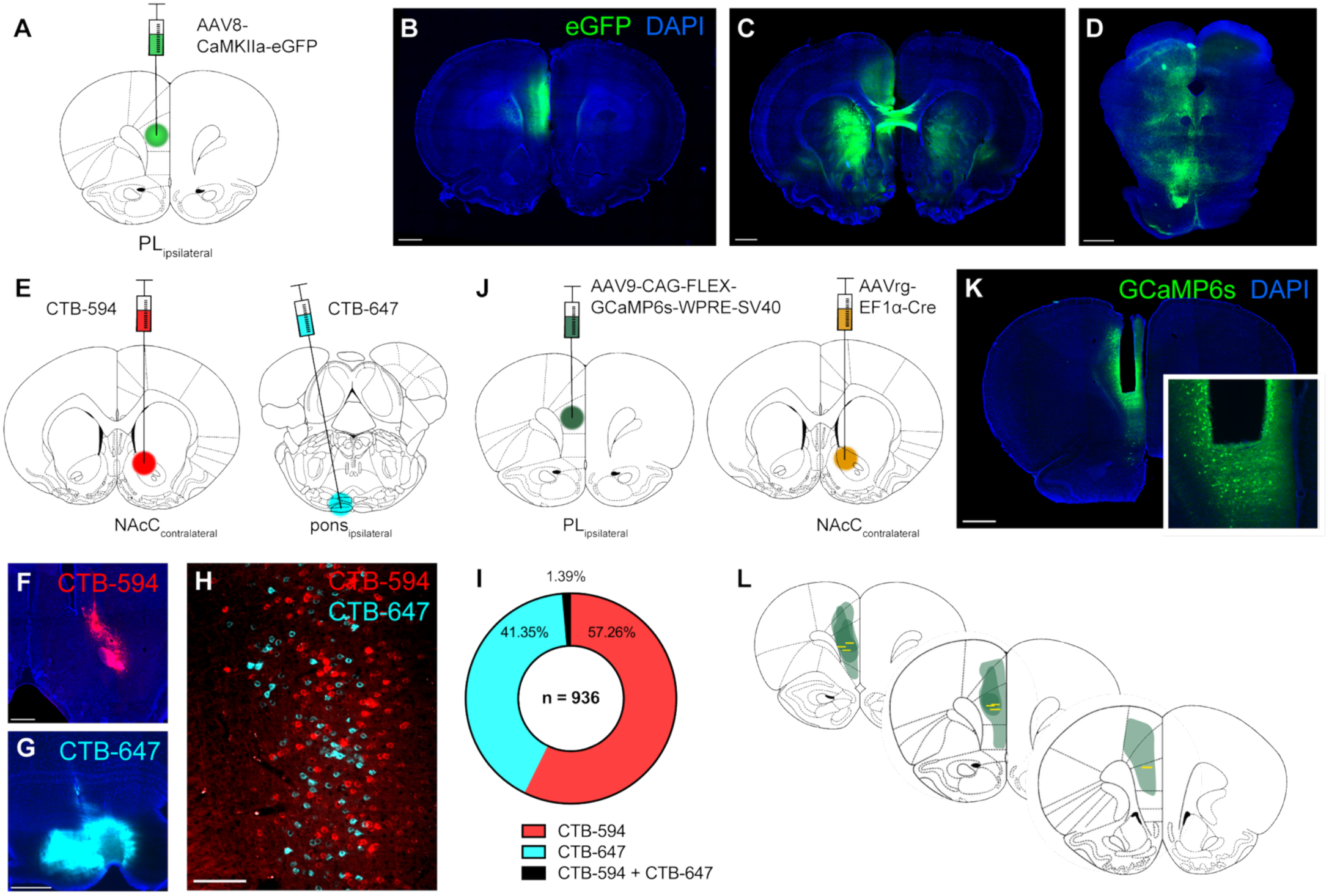
Intersectional virus strategy for cell-type selective GCaMP6s expression and fiber photometry recording from intratelencephalic (IT) neurons of the prelimbic cortex. **(A, B)** Injection of AAV8-CaMKIIa-eGFP into prelimbic cortex reveals axonal projections bilaterally throughout striatum and NAc **(C)** as well as hindbrain structures and the pyramidal tract **(D)**. Scale bars, 1000 μm. **(E – I)** Dual-retrograde tracing from contralateral NAcC and ipsilateral pons using fluorescent cholera toxin subunit B (CTB594, CTB-647) reveals distinct and nonoverlapping subpopulations of prelimbic projection neurons and demonstrates the efficacy of selective IT and PT targeting via downstream axonal projection targets. Scale bars, 600 μm for F, G, 150 μm for H. N = 936 cells from 3 rats. **(J)** Intersectional virus strategy in which a Cre-dependent GCaMP6s is infused into the ipsilateral prelimbic cortex (AAV8-CAG-FLEX-GCaMP6s-WPRE-SC40) and a retrogradely transported Cre recombinase (AAVrg-EF1α-Cre) is infused into the contralateral NAcC. **(K)** Representative expression of GCaMP6s and placement of an optic fiber within the prelimbic cortex. GCaMP6s fluorescence is amplified with immunostaining using an antibody against GFP (green) and nuclei are visualized using a DAPI mounting medium (blue). Inset, dense and bright expression of GCaMP6s is visible under the fiber tract. Scale bar, 1000 μm. **(L)** GCaMP6s expression (green shading) typically spanned the dorsomedial prefrontal cortex, and all fibers were positioned within the prelimbic region (the surface of each fiber is indicated with yellow bars). N = 7 rats.

## RESULTS

### An intermittent access model reveals distinct addiction-like phenotypes and sex differences in fentanyl self-administration

Rats were trained to self-administer fentanyl before undergoing IntA SA, a PR test, extinction, and a cue-induced reinstatement test (Figure 1A). To classify risk-phenotype, behavior from eight metrics: (1) acquisition, (2) infusion latency, (3) intake, (4) consistency, (5) seeking, (6) motivation, (7) extinction, and (8) reinstatement was normalized and combined into a cumulative z-score. This approached revealed two predominant phenotypes, low-risk (41%) and high-risk (49%) (Figure 1B). High-risk rats exhibited positive z-scores for a greater number of behavioral metrics compared to low-risk rats (Figure 1D). In addition, compared to low-risk rats, high-risk rats acquired self-administration behavior more rapidly (Figure 1E, P = 0.008, Welch’s t-test) and discriminated between the active and inactive lever on the first day of training (Supplemental Figure 1, P = 0.002, REML mixed-effects model), while low-risk rats did not exhibit such a preference until the fourth day (P = 0.02). High-risk rats earned more infusions (Figure 1G, P < 0.0001, Welch’s t-test) with a shorter latency (Figure 1F, P < 0.0001, Welch’s t-test), had more consistent infusion intake (Figure 1H, P < 0.0001, Welch’s t-test), and a greater number of active lever presses during drug unavailable (DU) periods (Figure 1I, P < 0.0001, Welch’s t test), the PR test (Figure 1J, P = 0.001, Welch’s t-test), extinction (Figure 1K, P < 0.0001, Welch’s t test) and during the cued reinstatement test (Figure 1L, P < 0.0001,Welch’s t-test). We observed correlations between many of the behavioral metrics (Figure 1C, including strong correlations between overall fentanyl intake, infusion latency, consistency of intake, and drug-seeking lever presses (Spearman’s Rho ranging from 0.50 to 0.93). Cue-induced reinstatement was significantly correlated with overall fentanyl intake (r = 0.43), infusion latency (r = 0.42), extinction lever presses (r = 0.61), consistency of intake (r = 0.38), and lever presses in the PR test (r = 0.58).

A central hallmark of addiction is escalation of intake, so we examined whether there were risk-phenotype and sex differences in this behavior. Overall, there was an increase in active lever presses (Figure 2A, B) and fentanyl infusions (Figure 2E, F) across the 13 days of IntA SA. Although low- and high-risk rats showed similar rates of escalation of active lever presses (Figure 2A) and fentanyl infusions (Figure 2E), high-risk rats had substantially higher daily presses (Figure 2A, main effect of severity, low- versus high-risk: F_(1, 44)_ = 68.26, P < 0.0001; main effect of time, days of IntA SA: F_(1, 43)_ = 10.38, P < 0.0001; two-way RM ANOVA) and levels of intake (Figure 2E, main effect of severity: F_(1, 44)_ = 76.32, P < 0.0001; main effect of time, days of IntA SA: F_(6.217, 267.3)_ = 11.37, P < 0.0001; two-way RM ANOVA). High-risk rats also showed greater active lever presses during the 4-second infusion and cue light timeout period (data not shown; main effect of severity, low- versus high-risk: F_(1, 45)_ = 17.06, P = 0.0002; daily average of 14 ± 1.00 timeout presses for high-risk, 4 ± 0.37 for low-risk), as well as greater inactive lever pressing during drug available (DA) periods (Figure 2A, main effect of severity, low- versus high-risk: F_(1, 44)_ = 5.54, P = 0.0231; main effect of time, days of IntA SA: F_(6.683, 287.4)_ = 3.82, P = 0.0007; two-way RM ANOVA).

To assess differences in drug-seeking behavior, we first examined lever presses during the IntA drug unavailable (DU) periods. During the first IntA session, rats had very high rates of active compared to inactive lever presses during DU periods (Figure 2C; high-risk, 114.64 ± 12.66 active lever presses, 29.76 ± 5.26 inactive lever presses; low-risk, 43.67 ± 6.52 active lever presses, 12.95 inactive lever presses). Although there was a steep rate of decline in unrewarded active lever presses as rats quickly learned to discriminate between DA and DU periods, active lever presses during DU periods remained significantly higher than inactive presses across sessions (Figure 2C; low-risk, main effect of lever: F_(1, 40)_ = 44.74, P < 0.0001; high-risk, main effect of lever: F_(1, 48)_ = 49.79, P < 0.0001). High-risk rats had a higher level of both active lever (Figure 2C, main effect of severity, low- versus high-risk: F_(1, 44)_ = 26.02, P < 0.0001; main effect of time, days of IntA SA: F_(12, 516)_ = 16.67, P < 0.0001; two-way RM ANOVA) and inactive lever (Figure 2C, main effect of severity, low- versus high-risk: F_(1, 44)_ = 20.65, P < 0.0001; main effect of time, days of IntA SA: F_(4.934, 212.2)_ = 5.99, P < 0.0001; two-way RM ANOVA) presses compared to low-risk rats. High-risk rats also exhibited greater DU active lever presses early in the IntA SA paradigm and thus showed a steeper rate of decline in pressing over the course of IntA SA (severity x time interaction, active lever: F_(12, 528)_ = 4.16, P < 0.0001).

To determine if there were sex differences in risk-phenotype and self-administration behaviors, we first examined the eight metrics in males and females. Females had higher cumulative z-scores (Figure 1B, P = 0.002, Welch’s t test), scored positive for more metrics and a greater proportion were classified as high-risk (Figure 1D, 75% for females versus 26% for males) compared to males. Females also had significantly greater levels of behavior on all metrics except for the latency to earn infusions and PR responding (Figure 1E-L P = 0.005-0.04, Welch’s t test). Examination of self-administration across IntA SA days revealed that, despite males and females showing similar rates of escalation, female rats had higher daily active lever presses (Figure 2B, main effect of sex: F_(1, 49)_ = 11.32, P = 0.002; main effect of time, days of IntA SA: F_(5.680, 278.3)_ = 13.22, P < 0.0001; two-way RM ANOVA), greater fentanyl intake (Figure 2F, main effect of sex: F_(1, 49)_ = 7.98, P = 0.007; main effect of time, days of IntA SA: F_(6.387, 312.9)_ = 13.80, P < 0.0001; two-way RM ANOVA) and more timeout period active lever presses (data not shown; main effect of sex, male vs female: F_(1, 50)_ = 8.53, P = 0.005; daily average of 13 ± 0.92 timeout presses for females, 6 ± 0.65 for males), and higher inactive lever presses (Figure 2B, main effect of sex: F_(1, 49)_ = 4.87, P = 0.032; main effect of time, days of IntA SA: F_(7.393, 362.2)_ = 4.72, P < 0.0001; two-way RM ANOVA) compared to male rats. Similarly, during DU periods female rats had a higher level of both active lever (Figure 2D, main effect of sex: F_(1, 49)_ = 4.67, P = 0.036; main effect of time, days of IntA SA: F_(4.532, 222)_ = 18.47, P < 0.0001; two-way RM ANOVA) and inactive lever (Figure 2D, main effect of sex: F_(1, 49)_ = 10.31, P = 0.002; main effect of time, days of IntA SA: F_(5.085, 249.2)_ = 6.56, P < 0.0001; two-way RM ANOVA) presses compared to male rats. Females also showed a high initial level of DU active lever pressing that sharply decreased over the first several days of IntA SA (sex x time interaction, active lever: F_(4.532, 222)_ = 4.34, P = 0.001).

### Representation of fentanyl-associated stimuli by intratelencephalic projection neurons of the prelimbic cortex across acquisition and escalation of fentanyl self-administration

To determine how IT neurons of the prelimbic cortex encode fentanyl-associated events including discriminative and conditioned cues, lever presses, and infusions, an intersectional viral strategy was used to express the genetically encoded calcium sensor GCaMP6s with cell-type specificity. We first used viral expression of eGFP to visualize prelimbic projections (Figure 3A, B), which revealed dense axonal collaterals throughout the striatum as well as hindbrain structures and the pyramidal tract (Figure 3C, D). We then validated our intersectional approach by infusing fluorophore-conjugated cholera toxin subunit B (CTB-594 and CTB-647) into IT- and PT-specific projection targets (Figure 3E). Retrograde tracing from contralateral nucleus accumbens core (NAcC_contralateral_) and ipsilateral pons revealed distinct and nonoverlapping subpopulations of prelimbic projection neurons, demonstrating the efficacy of selective IT – PT targeting via distinct downstream axonal projections (Figure 3F-I). Infusion of a Cre-dependent GCaMP6s AAV into the prelimbic cortex and a retrograde Cre vector into NAcC_contralateral_ (Figure 3J) produced dense GCaMP6s expression around optic fibers implanted in prelimbic cortex (Figure 3K). CGaMP6s expression typically spanned much of the dorsomedial PFC and the optic fiber was properly placed within prelimbic cortex in all rats (Figure 3L). Calcium signals were acquired using a wireless fiber photometry system that enabled long-term recordings from freely-moving rats in standard operant chambers across days of fentanyl self-administration (Figure 4A, B).

**Figure 4.**
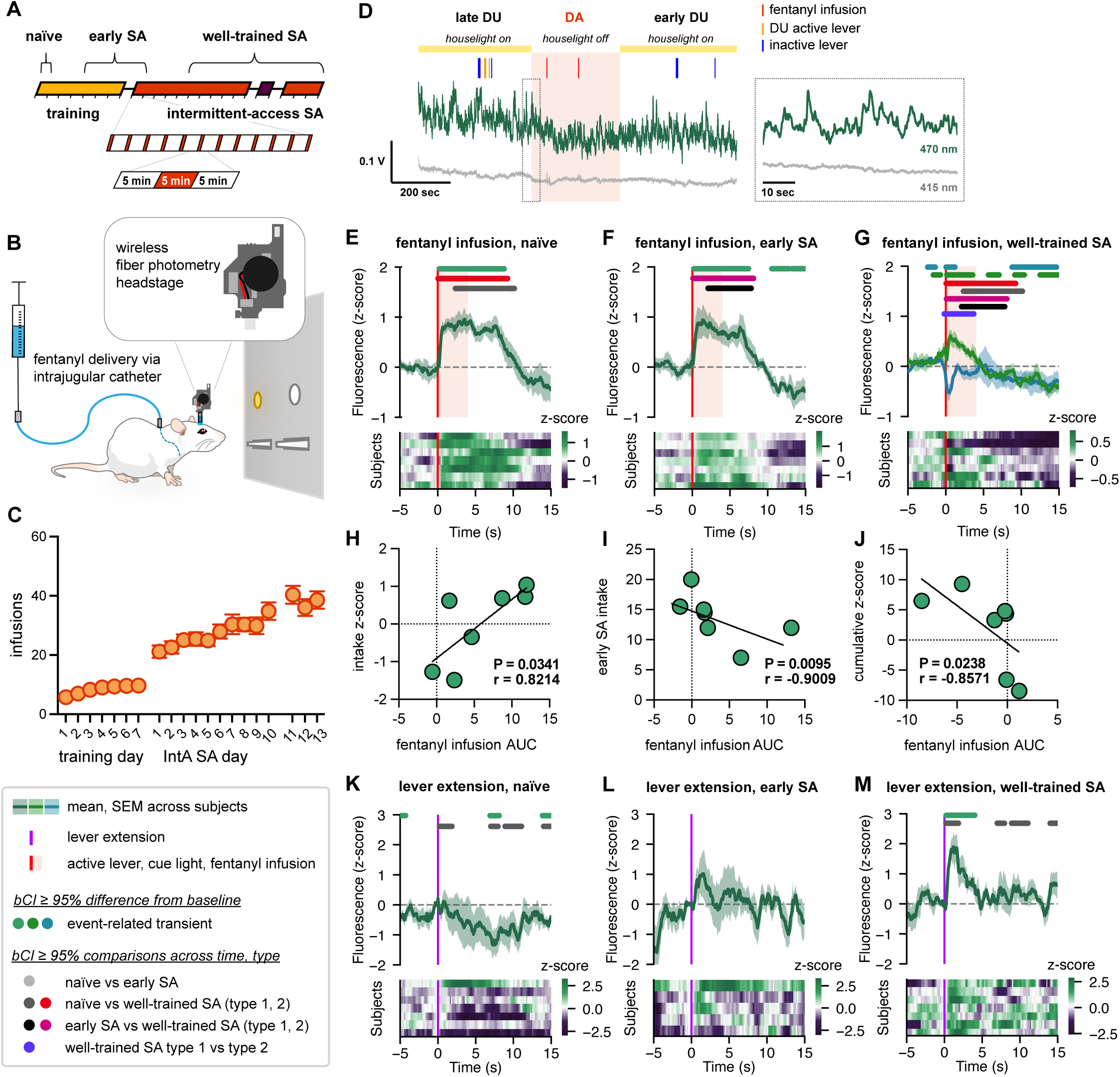
Prelimbic IT neurons exhibit dynamic encoding of fentanyl infusion, inactive lever presses, and DU active lever presses across acquisition and escalation of fentanyl SA. **(A)** Timeline of photometry recordings for lever presses and fentanyl infusions, in which recordings were performed on the first day of SA training (naïve), 1-2 days spanning the 5^th^ day of training to the first day of IntA SA (early SA), and 2-3 days spanning the 6^th^ day to the 13^th^ day of IntA SA (well-trained SA). **(B)** A wireless fiber photometry system was used to record IT calcium activity across days of fentanyl SA. **(C)** Rats gradually escalated fentanyl infusions across training and IntA SA days. **(D)** A representative IntA SA recording consisting of approximately 15 minutes spanning late DU, the entirety of DA, and early DU. Ticks are shown for inactive lever presses (blue), DU active lever presses (orange), and DA active lever presses (fentanyl infusions, red). GCaMP6s fluorescence (470 nm) is displayed in dark green and the isosbestic signal (415 nm) in gray. **(E – G, K – M).** Above, mean peri-event GCaMP6s fluorescence averaged across animals. Significant event-related transients (ERTs) and comparisons identified via bootstrapped confidence interval (bCI) are indicated with horizontal bars. Below, heatmaps of within-animal averaged peri-event fluorescence. Lower left, legend for photometry panels. N = 5 rats for lever extension in early SA, and 7 rats for all other data. **(E – G)** Fentanyl infusion ERTs diminished across naïve, early SA, and well-trained SA timepoints, and heterogenous response patterns are observed in well-trained SA. **(H)** In naïve rats the magnitude of the fentanyl infusion ERTs, measured as an area-under-the curve spanning the 15 seconds following lever press, positively correlated with the overall fentanyl intake z-score (used for severity classification and presented in Figure 1) for each animal. **(I)** In early SA, the magnitude of the fentanyl infusion ERTs negatively correlated with fentanyl intake during early SA sessions. **(J)** In well-trained rats, the magnitude of the fentanyl infusion ERTs negatively correlated with overall cumulative z-scores reflective of addiction severity. **(K – M)** IT calcium responses associated with initial lever extension at session onset increased in magnitude over the course of the SA paradigm.

We first examined prelimbic IT calcium activity associated with lever presses and fentanyl infusions over the course of SA, spanning 21 days across training and IntA SA sessions (Figure 4C, D). In recordings on the first day of SA training (naïve), there was an elevation in GCaMP6s fluorescence following active lever presses that was long-lasting relative to the fentanyl infusion (2.8 seconds) and cue light illumination (4 seconds), indicative of robust IT activation (Figure 4E). Notably, this reward-associated calcium response diminished in duration and amplitude across days of fentanyl SA, and bCI analysis (see methods) confirmed significant differences in ERTs recorded from well-trained rats compared to the earlier timepoints (Figure 4E-G). In recordings from early SA (days 5-8 of fentanyl SA), we observed a significant decrease in fluorescence following the initial elevation, occurring approximately 10 seconds after infusion onset (Figure 4F). By mid and late timepoints (well-trained SA, days 13-21) two distinct IT response patterns had emerged (Figure 4G). While a subset of animals continued to exhibit elevated calcium signals following rewarded active lever presses (designated type 1 responses), the remaining rats developed an inhibitory response with sharp decreases in fluorescence following these presses (type 2). Both response patterns were accompanied by decreases in fluorescence following fentanyl infusion and cue light illumination, as well as significant increases in fluorescence prior to the lever presses, while no such preceding increase was observed at earlier timepoints (Figure 4E, F). We confirmed the emergence of this approach-window calcium transient in well-trained SA by measuring the area under the curve (AUC) of the three seconds preceding the lever press, finding that fluorescence significantly differed from a baseline of 0 for well-trained SA (Figure 4G, P = 0.047; Wilcoxon test), but not for early SA (Figure 4F, P = 0.813) or naïve timepoints (Figure 4E, P = 0.578).

We also performed non-parametric resampling-based permutation tests to detect significance differences within specific windows across timepoints (Figure 4E-G)^69,70^. While no significance difference was found in comparisons of naïve and early SA timepoints of peri-event fluorescence in the 10 seconds following active lever presses (P = 0.535), we did detect differences between naïve signals and those from well-trained SA (naïve vs well-trained SA type 1, P = 0.005; naïve vs well-trained SA type 2, P = 0.01), as well as between early SA and type 2 responses in well-trained SA (P = 0.047). Additionally, type 1 and type 2 responses from well-trained rats differed significantly (type 1 vs type 2, P = 0.027, -3 to +4 seconds around lever press).

Noting heterogeneity in IT calcium responses across animals, we examined if IT activity patterns related to the severity of addiction-like behavior. We found significant correlations between fentanyl infusion ERTs and several behavioral metrics, including measurements reflective of self-administration behavior on recording days as well as the z-scored metrics used for severity classification (Figure 1). In naïve rats, the magnitude of the fentanyl infusion ERTs (measured as AUC following lever presses) positively correlated with overall fentanyl intake z-scores (Figure 4H, P = 0.034, r = 0.821). In contrast, during early SA, infusion ERT magnitude negatively correlated with fentanyl intake during early SA recording sessions (Figure 4I, P = 0.001, r = -0.901), as well as DA active lever press rate (Supplemental Figure 2A) and proportion of DA active lever presses occurring in bursts (≥3 presses with ≤10 second inter-press intervals; Supplemental Figure 2B). Additionally, early SA infusion ERT magnitude negatively correlated with overall latency and consistency z-scores (Supplemental Figure 2C, D). In well-trained rats, infusion ERTs negatively correlated with the cumulative z-score reflective of addiction severity (Figure 4J, P = 0.024, r = -0.857) as well as z-scores for motivation (Supplemental Figure 2I), seeking (Supplemental Figure 2J), and reinstatement (Supplemental Figure 2K).

Although these negative correlations initially seemed contradictory to our findings from naïve rats (Figure 4H), we noted that high- and low-risk rats exhibited diverging trajectories of infusion-associated calcium response across days of IntA SA. High-risk rats in our photometry cohort (5/7 rats) exhibited significantly diminished infusion ERTs by early SA (AUCs of 7.77 ± 2.01 in naïve recordings, 0.77 ± 0.69 in early SA), while in contrast low-risk rats (2/7) exhibited significantly larger ERTs in early SA (0.9068 ± 1.4508 in naïve recordings, 9.8894 ± 3.3459 in early SA). By well-trained timepoints, infusion ERTs from high-risk rats had further diminished, and responses from low-risk rats also decreased in magnitude. These changes are illustrated by the averaged peri-event GCaMP6s fluorescence of each animal (Figure 4E-G, heatmaps sorted by cumulative z-score in descending order such that each row corresponds consistently to the same rats across panels). Taken together, these trends suggest that rats which exhibit more severe addiction-like behavioral phenotypes also display a distinct pattern of IT responsiveness marked by initially high levels of fentanyl-associated activation that rapidly decreases over days of IntA SA.

Within the first several days of training, rats became very responsive to lever extension at the initiation of the session. Reasoning that prelimbic cortex may encode lever extension as a discriminative stimulus for fentanyl availability, we examined IT calcium transients evoked by lever extension across multiple SA timepoints (Figure 4A). On the first day of training, lever extension did not evoke a consistent IT response beyond brief intervals of delayed decreased fluorescence (Figure 4K). By early SA there was no significant positive or negative response to lever extension (Figure 4L). At later timepoints, however, (well-trained SA) lever extension prompted a significant ERT with a sharp increase in fluorescence (Figure 4M). Permutation tests and bCI comparisons across timepoints confirmed the appearance of a lever extension-associated IT response in well-trained SA (naïve vs well-trained, P = 0.039). In addition, we measured the AUC of the lever extension response and found a significant increase in fluorescence during well-trained (P = 0.016; Wilcoxon test), but not naïve (P = 0.297) or early SA timepoints (P = 0.813).

We wondered how IT responses may differ during unrewarded lever presses, either of the inactive lever (Supplemental Figure 3A) or of the active lever during DU periods (Supplemental Figure 3B). On the first day of training, inactive lever presses were associated with small and delayed increases in IT calcium activity (Supplemental Figure 3C). This inactive lever press ERT decreased across days of self-administration and no significant differences from baseline were detected in early SA (Supplemental Figure 3D). In well-trained rats, we instead observed decreases in IT calcium activity following inactive lever presses (Supplemental Figure 3E). Comparisons in bCI analysis indicated delayed intervals during which ERTs from well-trained SA and earlier timepoints differed, and permutation tests confirmed a significant difference in GCaMP6s fluorescence from naïve and well-trained timepoints in a delayed window following inactive lever presses (+5 to +15 seconds, P = 0.039).

Finally, we examined IT activity during unrewarded active lever presses in DU periods. In early SA, these presses evoked minimal ERTs (Supplemental Figure 3F), whereas in well-trained rats they produced a biphasic response, with an initial increase followed by a prolonged decrease in GCaMP6s fluorescence (Supplemental Figure 3G). Notably, in early SA, rewarded active lever presses during DA periods led to significant calcium responses that were absent during DU active lever presses (infusion versus DU active lever response, P = 0.0095; permutation test). By well-trained SA, however, shifts in response patterns to both DA and DU active lever presses resulted in comparable ERTs (DU active lever vs DA type 1 response, P = 0.540; vs DA type 2 response, P = 0.497; permutation test), beyond a brief interval following lever presses during which DU active lever responses significantly differed from type 2 infusion responses (0-2.2 sec, 95% bCI).

### High-risk rats rapidly acquire intermittent patterns of fentanyl-seeking and fentanyl-taking with robust responsiveness to the discriminative stimulus

Reasoning that rats exhibiting more severe addiction-like behavioral phenotypes may be more responsive to discriminative stimuli signaling fentanyl availability, we next analyzed the temporal organization of active lever pressing within the IntA SA sessions and across days of self-administration to determine if the development of intermittent fentanyl-taking and -seeking behaviors differed by risk-phenotype and sex. Measurement of active lever presses within 60 second bins across the entire 335 minutes of daily IntA SA illustrated a robust shift in active lever pressing away from unrewarded DU periods and into DA periods across 10 days of self-administration (Supplemental Figure 4).

We observed that by the end of the first IntA SA session, some rats began to focus active lever pressing within drug available (DA) periods, suggesting acquisition of the discriminative stimulus. We measured the proportion of total active lever presses occurring within DA periods and found that this proportion significantly increased across the first day of IntA SA for high-risk, but not low-risk, rats (Supplemental Figure 4A, B; first three DA and DU periods, excluding first DA, versus last three DA and DU periods; low-risk, from 12.97±2.58% to 21.55±5.57%, P = 0.188; high-risk, from 16.34±2.43 to 39.52±3.51%, P < 0.0001; paired t test), consistent with rapid acquisition of the intermittent-access task. By late IntA SA (data averaged across days 8-10), fentanyl-seeking has become highly patterned across session, with pronounced spikes in active lever press rate observed at the onset of each DA period and low press rates during the DU periods, such that a majority of active lever presses occurred during DA periods for both low- and high-risk rats (Supplemental Figure 4C, D; low-risk, 60.44%±13.19; high-risk, 66.47±13.29%). In both early and late IntA SA days, we find a significant interaction between addiction severity and time bin (early IntA SA, severity x 60-second bin interaction: F_(21.53, 861.2)_ = 1.870, P = 0.0096; late IntA SA, severity x 60-second bin interaction: F_(6.272, 276)_ = 11.23, P < 0.0001), suggesting that while both low- and high-risk rats develop an intermittent pattern of fentanyl-seeking and -taking, active lever pressing by high-risk rats exhibits a more prominent temporal structure.

We also measured the overall rates of active lever pressing during DA and DU periods and found clear patterns across time and addiction severity phenotypes (Figure 5A). A three-way ANOVA revealed significant main effects of availability (DA versus DU: F_(1,43)_ = 204.4, P < 0.0001) and addiction severity (low- versus high-risk: F_(1,44)_ = 90.19, P < 0.0001), as well as significant interactions between availability, time (early versus late IntA SA), and severity (availability x severity interaction: F_(1,44)_ = 55.60, P < 0.0001; availability x time interaction: F_(1,44)_ = 47.03, P < 0.0001; availability x severity x time interaction: F_(1,44)_ = 8.899, P = 0.0046; three-way RM ANOVA), suggesting that across days of IntA SA, lever pressing by high-risk rats is more significantly influenced by signaled fentanyl availability. Importantly, during early IntA SA high-risk but not low-risk rats showed higher rates of active lever pressing during DA periods (P = 0.0023) as compared to DU periods, suggesting rapid acquisition of IntA SA. In both early and late IntA SA, high-risk rats exhibited higher rates of active lever pressing in both DA and DU periods (early IntA SA, DA, P < 0.0001, DU, P = 0.0001; late IntA SA, DA, P < 0.0001, DU, P = 0.0178).

**Figure 5.**
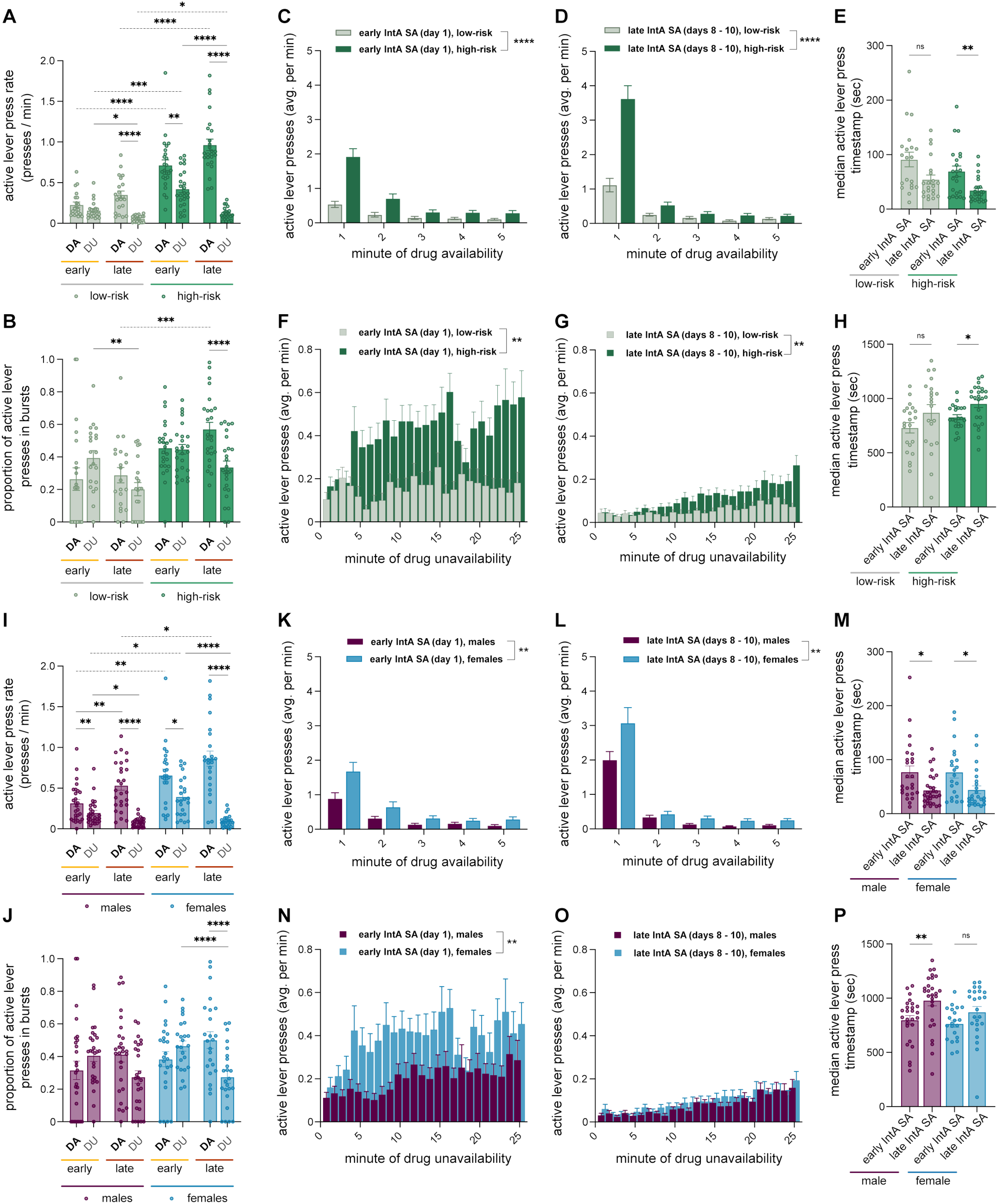
Temporal organization of fentanyl-taking and -seeking across early and late IntA SA. **(A)** Comparison of active lever press rates during DA and DU periods across early and late IntA SA revealed main effects and interactions of time, availability, and addiction severity phenotype. **(B)** Comparison of active lever press burst proportion during DA and DU periods across early and late IntA SA revealed main effects and interactions of time, availability, and addiction severity phenotype. **(C, D)** High-risk rats pressed the active lever at higher rates and with greater temporal structure across DA periods than low-risk rats. **(E)** From early to late IntA SA high-risk rats, but not low-risk, shifted lever presses to earlier within DA periods. **(F, G)** High-risk rats exhibited a higher rate of seeking lever presses than low-risk rats and escalated seeking within DU periods in both early and late IntA SA. **(H)** The timing of seeking lever presses shifted to later in DU periods from early to late IntA SA for high-risk rats. **(I)** Comparison of active lever press rates during DA and DU periods across early and late IntA SA revealed main effects and interactions of time, availability, and sex. **(J)** Male and female rats exhibited similar changes in active lever press burst proportions during DA and DU periods across early and late IntA SA, although only females showed greater burst pressing during DA of late IntA SA. **(K, L)** Although females pressed the active lever at higher rates during DA periods, the temporal structure of lever pressing was similar across sexes in late IntA SA. **(M)** From early to late IntA SA both males and females shifted lever presses to earlier within DA periods. **(N, O)** Female rats exhibited a higher rate of seeking lever presses than males in early IntA SA, but no sex differences were found in late IntA SA, and rates of escalation were similar. **(P)** The timing of seeking lever presses shifted to later in DU periods from early to late IntA SA for male rats.

We reasoned that highly motivated fentanyl-taking may appear as rapid successive active lever presses and therefore measured the proportion of active lever presses occurring in bursts (minimum of three presses, maximum inter-press interval of 10 seconds). High-risk rats showed a greater burst pressing compared to low-risk rats (Figure 5B; main effect of addiction severity F_(1,44)_ = 17.34, P = 0.0001; availability x severity interaction: F_(1,44)_ = 9.714, P = 0.0032; availability x time interaction: F_(1,43)_ = 29.80, P < 0.0001; three-way RM ANOVA). Notably, high-risk but not low-risk rats showed a higher burst proportion in DA periods compared to DU periods during late IntA SA (P < 0.0001). During late IntA SA high-risk rats exhibited a greater proportion of active lever presses in bursts in DA (P = 0.0007), but not DU periods, as compared to low-risk rats.

We hypothesized that rats exhibiting more addiction-like behavioral phenotypes may show more responsiveness to DA onset as well as distinct patterns of fentanyl-seeking during DU periods as compared to low-risk rats. We therefore averaged lever press rates in 60-second bins across DA and DU periods to examine the temporal organization of active lever pressing in early and late IntA SA. In both early and late IntA SA, high-risk rats exhibited a higher rate of active lever pressing in DA periods with a more pronounced temporal structure, such that most lever pressing occurred soon after the onset of availability (Figure 5C, early IntA SA, main effect of severity: F_(1,40)_ = 31.67, P < 0.0001; main effect of time, 1 minute bins: F_(1.758, 68.58)_ = 47.09, P < 0.0001; severity x time bin: F_(1.714, 68.54)_ = 14.58, P < 0.0001; Figure 5D, late IntA SA, main effect of severity, F_(1,43)_ = 51.26, P < 0.0001; main effect of time, 1 minute bins: F_(1.112, 46.69)_ = 90.44, P < 0.0001; severity x time bin interaction: F_(1.113, 47.84)_ = 24.56, P < 0.0001; two-way RM ANOVA). Measurement of active lever press timestamps within DA periods indicated that high-risk rats, but not low-risk rats, shifted active lever pressing to earlier timepoints across days of IntA SA, consistent with greater responsiveness to the discriminative stimulus signaling fentanyl availability (Figure 5E; low-risk, early vs late IntA SA, P = 0.095; high-risk, early vs late IntA SA, P = 0.005; Kruskal-Wallis test).

Although low-risk and high-risk rats showed comparable active lever pressing in the first several minutes of the DU periods, high-risk rats pressed at increasingly higher rates as compared to their low-risk counterparts resulting in a clear escalation of fentanyl-seeking by high-risk rats over the course of the DU periods in both early IntA SA (Figure 5F; main effect of severity, F_(1,40)_ = 19.38, P < 0.0001; main effect of time, 1 minute bins: F_(10.80, 421)_ = 2.483, P = 0.005; severity x time bin interaction: F_(10.99, 439.5)_ = 1.92, P = 0.035; two-way RM ANOVA) and late IntA SA (Figure 5G; main effect of severity: F_(1, 44)_ = 10.67, P = 0.002; severity x time bin interaction, F_(7.308, 321.6)_ = 2.740, P = 0.008). Although both phenotypes exhibited escalation of fentanyl-seeking during DU periods in late IntA SA, high-risk rats escalated their lever presses with a significantly steeper slope than low-risk rats (early IntA SA, P = 0.0015; late IntA SA, P < 0.0001; simple linear regression). From early to late IntA SA active lever pressing occurred later in DU periods for high-risk, but not low-risk rats, further supporting increased escalation of fentanyl-seeking across days of IntA SA (Figure 5H; low-risk, early vs late IntA SA, P = 0.058; high-risk, early vs late IntA SA, P = 0.037; Kruskal-Wallis test).

Altogether these data reveal that, although shifts in behavioral patterns over the course of IntA SA are observed in both low- and high-risk groups, high-risk rats exhibit particularly pronounced acquisition and refinement of fentanyl-taking and fentanyl-seeking within the intermittent task structure. High-risk rats exhibit high rates of active lever pressing preferentially within DA periods early in the IntA SA paradigm, and bursts of active lever pressing during DA periods in late IntA SA. High-risk rats also earn fentanyl infusions increasingly rapidly in the signaled periods of drug availability across days of IntA SA, and exhibit marked escalation of seeking during DU periods, consistent with robust responsiveness to the discriminative stimulus and highly motivated fentanyl-taking and -seeking.

In light of our prior observations that female rats predominantly displayed a high-risk phenotype, we wondered whether we would find a sex difference in the temporal organization of lever pressing in early or late IntA SA. Although we found overall higher rates of active lever pressing by female rats (Supplemental Figure 4; Figure 5I; main effect of sex, F_(1,49)_ = 18.85, P < 0.0001), consistent with higher daily lever press totals (Figure 2), we did not observe earlier acquisition of the discriminative stimulus by females. We found that the proportion of total active lever presses occurring in DA periods increased across the first day of IntA SA for both male and female rats (Supplemental Figure 4E, F; male, from 14.05±1.99% to 30.51±5.20%, P = 0.003; female, from 15.37±2.72 to 36.18±4.85%, P = 0.002; paired t test), and both males and females exhibited higher rates of lever pressing in DA periods as compared to DU periods across early and late IntA SA (Figure 5I; male, early: P = 0.0032; female, early: P = 0.0147; male, late: P < 0.0001; female, late: P < 0.0001). However, while in early IntA SA females showed a higher rate of lever pressing than males regardless of fentanyl availability (DA, P = 0.0081, DU, P = 0.0161), in late IntA SA female rats showed higher rates of lever pressing specifically during DA periods (P = 0.0487), and analysis of binned lever presses across the 335 min sessions of late IntA SA revealed a sex x time bin interaction (F_(4.620, 226.4)_ = 2.811, P = 0.0204; two-way RM ANOVA). Similarly, our two-way RM ANOVA of lever press rates revealed a sex x availability interaction (F_(1,49)_ = 12.44, P = 0.0009). In late IntA SA, females but not males exhibited a greater proportion of active lever press bursts in DA periods as compared to DU periods (Figure 5J, P < 0.0001).

We compared the temporal organization of active lever presses during DA periods and found broadly similar patterns across time for males and females, with increased lever press rates in late IntA SA primarily within the first minute of DA periods (Figure 5K, L) and comparable shifts in lever press timestamps (Figure 5M, male, early vs late IntA SA, P = 0.0201; female, early vs late IntA SA, P = 0.0145; Kruskal-Wallis test). Although we detected a higher rate of lever pressing in DA bins by females (main effect of sex: early IntA SA, F_(1,45)_ = 11.26, P = 0.0016; late IntA SA, F_(1,49)_ = 9.682, P = 0.0031), we found a sex x bin interaction only in early IntA SA (F_(1.52, 68.30)_ = 3.85, P = 0.037; two-way RM ANOVA), suggesting no sex difference in the temporal pattern of fentanyl-taking in late IntA SA. Although females displayed a higher rate of seeking lever presses during DU periods of early IntA SA (Figure 5N; F_(1,45)_ = 9.87, P = 0.003), we observed no sex difference in DU pressing in late IntA SA (Figure 5O; F_(1,49)_ = 0.9441, P = 0.336), and we found a comparable escalation of fentanyl-seeking across the DU periods in both early and late IntA SA (early IntA SA, P = 0.9001, late IntA SA, P = 0.8580, simple linear regression).

Altogether, although we find higher rates of lever pressing during fentanyl availability by females, reflective of greater fentanyl intake across the IntA SA paradigm, we observe a comparable temporal structure to fentanyl-seeking and -taking across sexes, suggesting similar acquisition of the discriminative stimulus by males and females.

### Prelimbic IT neurons encode discriminative and conditioned cues signaling fentanyl availability and reward

Given the patterns of binge-like fentanyl-taking and -seeking, we examined how prelimbic IT neurons encode the conditioned and discriminative cues that sculpt addiction-like behavior across IntA SA. Drug availability (DA) was signaled by extinguishing of the houselight and prelimbic IT neurons were highly responsive to this cue, with significant elevations in GCaMP6s fluorescence observed during early (Figure 6B), mid (Figure 6C), and late (Figure 6D) IntA SA sessions. Although bCI analysis revealed a decreased duration of the DA cue-evoked response over time, with significant decreases in the descending phase of the calcium transient, the mean peak amplitude of the ERT remained stable. Transitions to DU periods, signaled by illumination of the houselight, similarly elicited significant IT neuron responses at all timepoints (Figure 6H, I, J). However, this DU cue-evoked response tended to not diminish, with only brief decreases in fluorescence from mid to late IntA SA (Figure 7J, K). Direct comparison of cue responses revealed that IT activity was greater for DA versus DU cue onset during early SA (P = 0.0245, permutation test), but were not different at mid (P = 0.3525) or late sessions (P = 0.5615).

**Figure 6.**
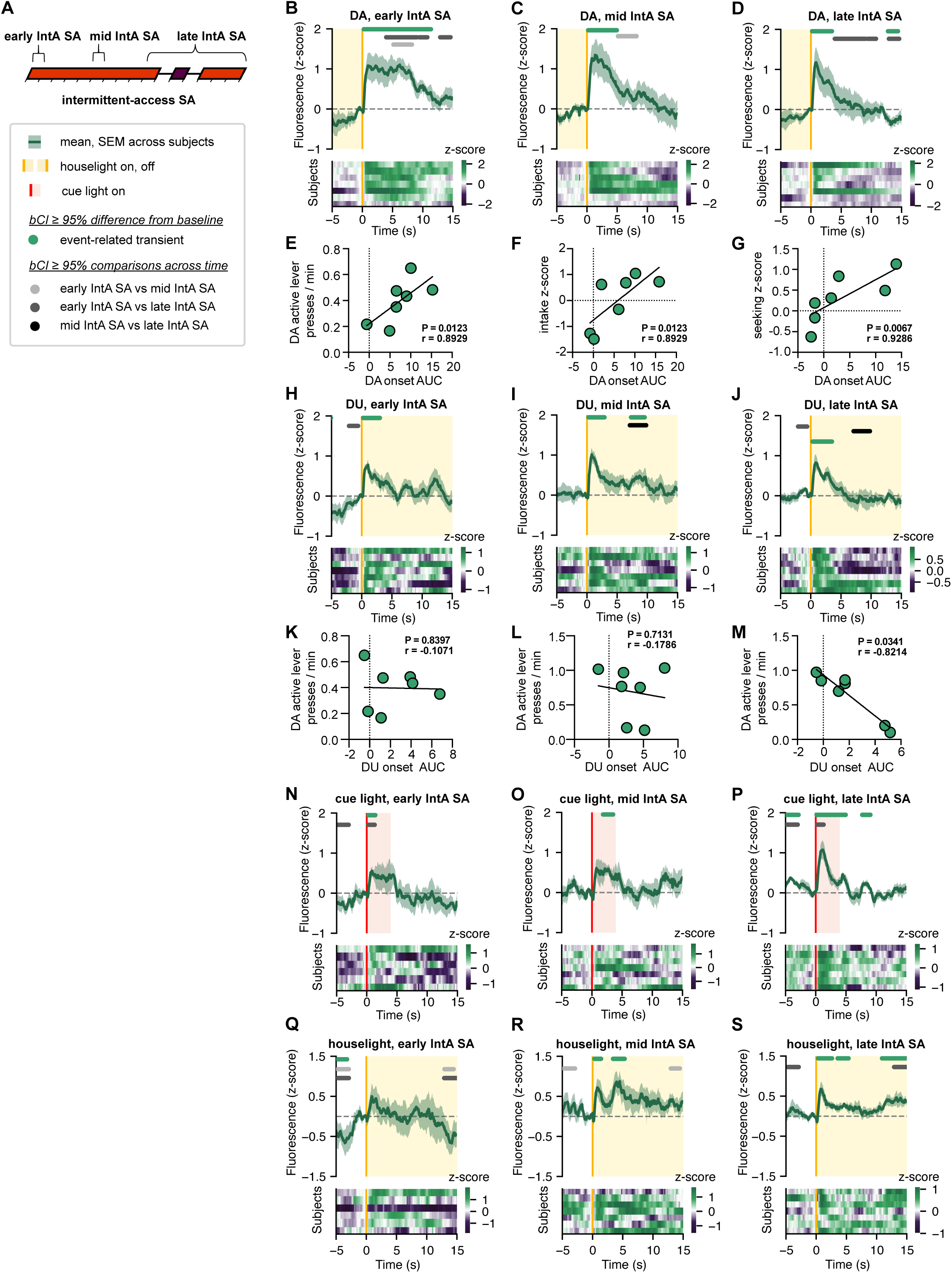
Prelimbic IT neurons develop robust responses to conditioned and discriminative stimuli associated with fentanyl reward and availability over the course of fentanyl SA. **(A)** IT responses to onset of drug availability and unavailability were recorded on the first day of IntA SA (early), the 6^th^ day of IntA SA (mid), and 2-3 days spanning the 10^th^ day to the 13^th^ day of IntA SA (late). Below, legend for panels B – D, H – J, N - S. Above, mean peri-event GCaMP6s fluorescence averaged across animals. Significant event-related transients (ERTs) and comparisons identified via bootstrapped confidence interval (bCI) are indicated with horizontal bars. Below, heatmaps of within-animal averaged peri-event fluorescence. N=7 rats for all panels. **(B – D)** ERTs associated with DA onset decreased in duration but remained stable in amplitude over the course of IntA SA. **(E)** In early IntA SA, ERTs associated with DA onset positively correlated with the rate of DA active lever presses. **(F)** In mid IntA SA, DA onset ERTs positively correlated with the overall fentanyl intake z-score. **(G)** In late IntA SA, DA onset ERTs positively correlated with the seeking z-score. **(H – J)** ERTs associated with DU onset were characterized by a relatively stable increase in fluorescence. **(K – M)** The ERT associated with DU onset negatively correlated with the rate of DA active lever presses in late IntA SA, but not earlier timepoints. **(N – P)** IT response to non-contingent conditioned cue presentation increased in duration and magnitude over the course of IntA SA. **(Q – S)** Non-contingent presentation of the discriminative houselight cue that signaled drug unavailability did not produce an ERT in early IntA SA, while a significant response was found in mid and late IntA SA.

**Figure 7:**
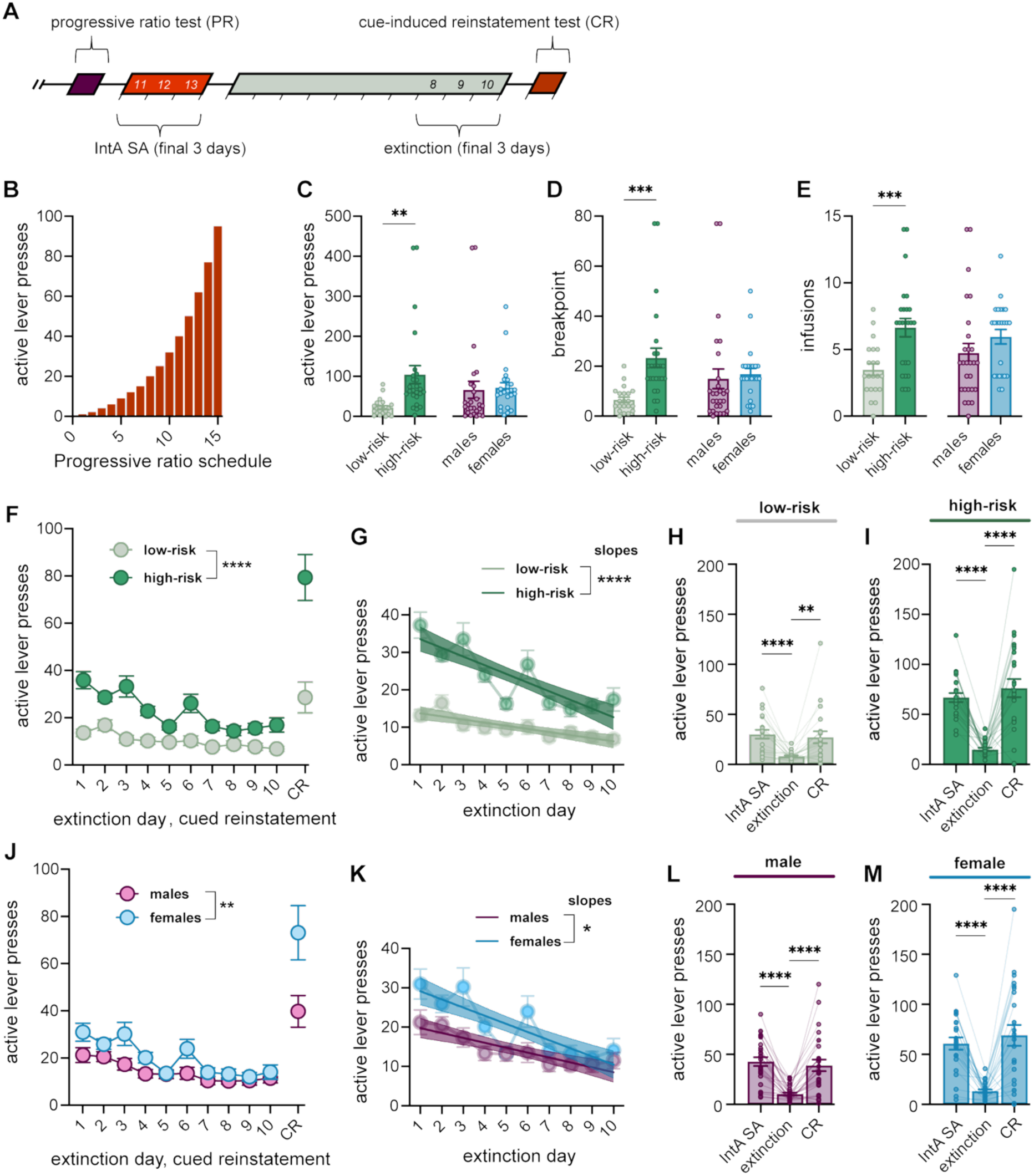
Sex and individual differences in fentanyl-taking and fentanyl-seeking during the progressive ratio test, extinction training, and cue-induced reinstatement test. **(A)** Timeline of progressive ratio testing (PR), IntA SA re-baseline, extinction training, and the cue-induced reinstatement test (CR). Representative IntA SA and extinction daily active lever press totals are averages from days 11 – 13 of IntA SA and days 8 – 10 of extinction training, while the CR active lever press total is measured from one 60 minute test that occurs on the day following 10 days of extinction training. **(B)** The progressive ratio schedule of reinforcement. **(C – E)** High-risk rats exhibit greater active lever pressing, a higher breakpoint, and earn more infusions than low-risk animals. **(F, G)** High-risk rats exhibited distinctly higher levels of extinction lever pressing than low-risk rats, with both a higher total number of active lever presses and a more prominent decline in pressing over the course of extinction training. **(H, I)**. Both low-risk and high-risk animals significantly extinguished active lever pressing and subsequently reinstated active lever pressing upon cue presentation. **(J, K)** Female rats exhibited greater extinction lever pressing than male rats, with both a higher total number of active lever presses and a more prominent decline in pressing over the course of extinction training. **(L, M)** Both male and female animals significantly extinguished active lever pressing and subsequently reinstated active lever pressing upon cue presentation.

Notably, the IT calcium responses associated with DA and DU period onsets were significantly correlated with multiple measures of addiction-like behavior. During early IntA SA, ERTs occurring at DA onset were positively correlated with drug-taking behaviors, including DA active lever presses (Figure 6E, P = 0.01, r = 0.893), first day (Supplemental Figure 2E) and overall fentanyl intake, infusion latency and consistency (Supplemental Figure 2F – H). At the mid IntA SA timepoint, ERTs at DA period onset retained a positive correlation with overall fentanyl intake (Figure 6F, P = 0.01, r = 0.893), while by late IntA SA these events were instead correlated with drug-seeking (Figure 6G, P = 0.007, r = 0.929) and motivation (Supplemental Figure 2L). In contrast, IT calcium responses associated with the onset of DU periods did not significantly correlate with behavioral metrics until late IntA SA. Unlike DA onset signals, DU onset ERTs negatively correlated with drug-taking measures including DA active lever presses (Figure 6M, P = 0.034, r = -0.821) and fentanyl intake in late IntA SA (Supplemental Figure 2N) as well as drug-seeking measures such as DU active lever presses (Supplemental Figure 2M) and active lever presses during extinction (Supplemental Figure 2O).

Finally, we examined IT responses to non-contingent presentations of the conditioned stimulus (cue light) and discriminative stimulus (houselight) across IntA SA timepoints. We found significant cue light ERTs at all time points, and the duration of these transients increased over time (Figure 6N, O, P), as did their magnitude (bCI analysis). Non-contingent presentation of the houselight did not produce a detectable ERT until the mid IntA SA probe session, which appeared stable in magnitude thereafter (Figure 6Q, R, S). Measurement of the AUC of the cue light and houselight responses indicated significant fluorescence above baseline in late IntA SA (cue light, P = 0.0312; houselight, P = 0.0156; Wilcoxon test), but not earlier timepoints.

Altogether, these data reveal dynamic representation of fentanyl-associated stimuli by prelimbic IT neurons, such that by late IntA SA robust event-related calcium transients are observed to houselight-signaled drug availability, drug unavailability, and non-contingent presentations of both conditioned and discriminative stimuli. IT calcium transients associated with DA and DU period onsets appear to reflect ongoing self-administration (for example, correlating with active lever pressing on the day of recording) as well as predict subsequent addiction-like behavior (z-scored severity metrics including fentanyl-seeking during extinction and overall consistency of fentanyl intake across IntA SA).

### High-risk rats exhibit greater fentanyl intake during progressive ratio tests and greater fentanyl-seeking during extinction training and cued reinstatement

People with substance use disorders show high levels of motivation for drugs and drug-seeking, so we examined these behaviors in our model. During the PR test (Figure 7B) high-risk rats had greater active lever presses (Figure 7C, P = 0.001, Welch’s t test), a higher breakpoint (Figure 7D, P = 0.0002, Welch’s t test), and more infusions (Figure 7E, P = 0.0004, Welch’s t test) as compared to low-risk rats, suggesting greater levels of motivation for fentanyl.

Although both low- and high-risk rats showed a significant decline in active lever presses over the course of extinction training (Figure 7F-I, IntA SA versus extinction: low-risk, P < 0.0001; high-risk, P < 0.0001; paired t test) and a cue-induced increase in presses during the reinstatement test, indicating that presentation of the conditioned cue light was sufficient to restore fentanyl-seeking (Figure 7F-I, extinction versus CR: low-risk, P = 0.0028; high-risk, P < 0.0001; paired t test), high-risk rats had higher levels of extinction and reinstatement active lever presses than low-risk rats (Figure 7F, main effect of severity: F_(1, 44)_ = 51.58, P < 0.0001; main effect of time, days of extinction and CR: F_(2.298, 99.75)_ = 32.73, P < 0.0001; severity x time interaction: F_(2.274, 98.70)_ = 8.129, P = 0.0003; REML mixed-effects model) indicating greater fentanyl-seeking in the absence of drug reinforcement. Greater fentanyl-seeking by high-risk rats resulted in a steeper decline of active lever pressing across days of extinction training (Figure 7G, P < 0.0001, simple linear regression). Comparison of active lever pressing during the CR test confirmed greater reactivity to the presentation of the fentanyl-associated conditioned cue by high-risk rats (high-risk, 77.48 ± 9.070 presses, low-risk, 27.29 ± 5.968 presses, P < 0.0001, t test).

In contrast to the effects of risk-phenotype, there were no sex differences in the motivation for fentanyl (Figure 7C-E). However, although both males and females decreased active lever presses during extinction and increased presses during the reinstatement test (Figure 7J-M, IntA SA versus extinction: male, P < 0.0001; female, P < 0.0001; paired t test; Figure 7J-M, extinction versus CR: male, P < 0.0001; female, P < 0.0001; paired t test), females had higher levels of drug-seeking, as evidenced by more active lever presses during extinction and reinstatement (Figure 7J, main effect of sex: F_(1, 49)_ = 8.628, P = 0.0050; main effect of time, days of extinction and CR: F_(10, 480)_ = 33.19, P < 0.0001; sex x time interaction: F_(10, 480)_ = 4.191, P < 0.0001; REML mixed-effects model), and a steeper decline in active lever pressing across extinction (Figure 7K, P = 0.0397, simple linear regression). Our data reveal a more robust cue-induced reinstatement of fentanyl-seeking by female rats as compared to male rats (females, 69.04 ± 10.41 presses, males, 39.11 ± 5.763 presses, P < 0.0001, t test), despite a similar degree of active lever pressing by the end of the extinction training phase.

### Prelimbic IT calcium responses diminish across extinction and re-emerge during reinstatement of fentanyl-seeking induced by a conditioned cue

Prelimbic IT neurons become responsive to lever extension at the initiation of SA sessions (Figure 4), so we first examined whether this activity persisted across extinction training as rats decreased fentanyl-seeking (Figure 8A-C). The increases in GCaMP6s fluorescence upon lever extension were present during early and mid-extinction (Figure 8D, E). In late extinction, although there was still a significant increase in GCaMP6s fluorescence, the ERT was shorter in duration (Figure 8F).

**Figure 8.**
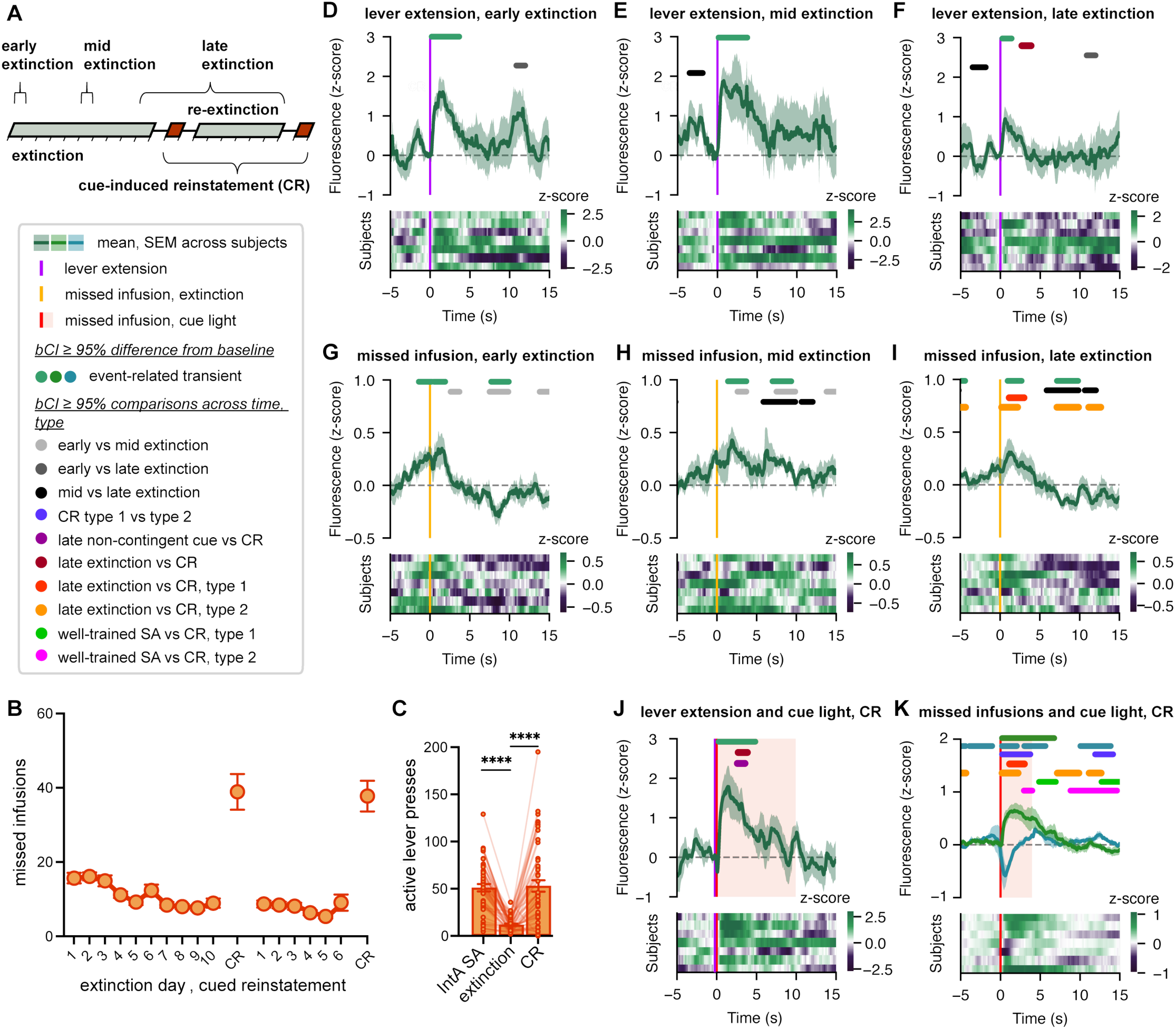
Prelimbic IT encoding of fentanyl-associated stimuli diminishes across extinction and re-emerges during cue-induced reinstatement. **(A)** Extinction ERTs were recorded on the first day of extinction training (early), the 6^th^ day of extinction (mid), and the 10^th^ day of extinction along with the 6^th^ day of re-extinction (late). CR photometry data was recorded from two sessions separated by 6 days of re-extinction. Below, legend for all photometry panels. **(B, C)** Fentanyl-seeking, measured as the number of daily missed infusions (unrewarded active lever presses), gradually decreased across extinction training but is reinstated by presentation of the conditioned stimulus. **(D – K)** Above, mean peri-event GCaMP6s fluorescence averaged across animals. Significant event-related transients (ERTs) and comparisons identified via bootstrapped confidence interval (bCI) are indicated with horizontal bars. Below, heatmaps of within-animal averaged peri-event fluorescence. N = 7 rats for all panels. **(D – F)** Lever extension ERTs were present at all timepoints but decreased in duration in late extinction. **(G – I)** ERTs associated with missed infusions included elevations in GCaMP6s fluorescence prior to lever presses in early, but not mid or late, extinction. **(J)** Conditioned cue light presentation accompanied lever extension in CR sessions and prompted a greater increase in GCaMP6s fluorescence than that observed during late extinction session onset or prior non-contingent cue presentations. **(K)** Two distinct types of ERTs were observed with CR missed infusions (active lever presses rewarded with cue illumination but no fentanyl delivery), reminiscent of those seen during late IntA SA, both of which differed significantly from late extinction missed infusion ERTs.

We next examined IT calcium activity associated with active lever presses that were previously followed by cue light illumination and fentanyl delivery (missed infusions, Figure 8B). On the first day of extinction, we observed increased GCaMP6s fluorescence just prior to and after lever presses (Figure 8G), similar to the increases during late IntA SA (Figure 4G). The elevation in fluorescence preceding lever presses was no longer different from baseline in mid and late extinction (Figure 8H, I), but there was a brief elevation of GCaMP6s that quickly declined after several seconds, leading to a sub-baseline decrease in fluorescence during the missed infusion period (Figure 8I).

During the cue-induced reinstatement test, initial lever extension was accompanied by a 10-second non-contingent presentation of the conditioned cue light previously paired with fentanyl infusions, which we observed was sufficient to induce robust reinstatement of fentanyl-seeking (from an average of 51.33 ± 3.846 daily presses across the final three days of IntA SA, to 11.96 ± 1.065 average daily presses across the final three days of extinction, and finally 53.20 ± 6.090 presses in the CR test; Figure 8C). This cue presentation resulted in a sharp and significant increase in GCaMP6s fluorescence (Figure 8J). The calcium response was greater than the response elicited by the lever extension in late extinction (Figure 8F, J) or by non-contingent cue presentations during IntA SA (Figure 6P, 8J).

When we examined IT GCaMP6s activity associated with subsequent active lever presses in the CR test, which were followed by cue light presentation but no fentanyl delivery (missed infusions), we observed heterogenous response patterns reminiscent of those seen during mid and late IntA SA. While most rats exhibited a clear increase in GCaMP6s fluorescence during missed infusions (type 1), a subset of animals exhibited a sharp decrease in fluorescence (type 2), which in these sessions was followed by a brief elevation in fluorescence (Figure 8K). Notably, of the three rats that previously exhibited type 2 responses in IntA SA, two once again exhibited a type 2 response in CR while the third instead exhibited a type 1 response, indicating that the response types observed in individual animals are not necessarily stable over time. Our bCI analysis revealed intervals of significant difference between late extinction ERTs and both type 1 and type 2 missed infusion ERTs in reinstatement sessions (Figure 8I, K). Taken together, these data indicate that cue light presentation during CR sessions substantially altered prelimbic IT calcium activity associated with both session onset and active lever presses.

## DISCUSSION

Our intermittent-access fentanyl self-administration paradigm recapitulates several features of addiction, including escalation of intake and persistent drug-seeking. We found that high-risk rats rapidly acquired an intermittent pattern of fentanyl-seeking and robust responsiveness to both conditioned and discriminative fentanyl-associated cues, as evidenced by binge-like bursts of fentanyl intake rapidly upon signaled drug availability and cue-induced reinstatement of fentanyl-seeking following extinction training. An intermittent pattern of drug intake is thought to more accurately recapitulate the pattern of drug use in humans and has been found to produce repeated “spikes” in drug concentration within the brain^20,72^, and evidence from animal models of addiction suggest that this pattern of drug intake has important implications for individual variation and sex differences in drug-seeking behaviors^20,73,74^. Our fentanyl IntA SA paradigm produces a subset of rats exhibiting an addiction-like behavioral phenotype, similar to that we’ve previously described in cocaine^63^ and heroin models^64^, and thereby provides a powerful and translationally-relevant model in which to pursue a mechanistic understanding of the genetic and neurobiological factors that underlie individual vulnerability to maladaptive substance use^12,13,44,71,75^.

Recognizing the accumulating evidence for important differences in male and female experience of substance use disorders, and the historical exclusion of female animals from many biological studies, investigations in rodent models of addiction have increasingly considered sex as an important variable in experimental design^16–19^. Considering reports of more severe experience of withdrawal in women opioid users^14^ along with greater cue-induced opioid-craving^15^, in the present fentanyl self-administration study we sought to characterize sex differences in addiction-like behaviors and describe how low- and high-risk behavioral phenotypes mapped onto male and female groups. We found that, compared to males, female rats were much more likely to be classified as high-risk and displayed more severe addiction-like behaviors across many of the metrics used for classification, including greater fentanyl intake and higher levels of fentanyl seeking during DU periods, extinction, and cued reinstatement.

Our findings contribute to a growing body of work describing sex differences in rodent models of opioid self-administration and echo several recent findings. A higher level of responding during cued reinstatement by females was recently reported in a study of continuous-access oral fentanyl SA^71^, as well as a model of heroin IntA SA^20^. In another study of 24-hour intermittent access to fentanyl, female rats earned a greater number of fentanyl infusions across more trials than males^19^, similar to our present findings of greater intake and consistency by females. However, we also note several discrepancies with past findings from continuous-access models. In continuous-access oral fentanyl SA, female rats did not exhibit higher levels of fentanyl intake than males, and male rats exhibited a greater number of active lever presses^71^, while in an 6-hour extended continuous-access paradigm authors reported greater reinstatement pressing in male rats, not females^18^. These differences may reflect the degree to which different temporal patterns of drug access and intake can uncover important nuances in addiction-like behaviors across individuals and sexes, as supported by numerous studies that have directly compared continuous and IntA paradigms^20,63,73^.

Longitudinal fiber photometry recordings revealed that prefrontal cortex IT neuronal activity dynamically tracks fentanyl-associated stimuli across the progression of drug use, implicating these cells in encoding aspects of addiction-like behavior. We found that the prelimbic IT population exhibited robust and consistent calcium responses to discriminative stimuli (lever extension and houselight-signaled fentanyl availability) that were often greater than those associated with conditioned cues or lever presses. These patterns of calcium activity support the notion that prelimbic cortex is highly attuned to operant contingencies and drug availability, not simply instrumental actions important for drug-taking, consistent with the established role for prelimbic cortex in regulating behavior according to discriminative stimuli. Pharmacological inactivation of prelimbic cortex was previously found to degrade performance in cocaine self-administration tasks involving periods of signaled availability and unavailability, indicated that prelimbic cortex functions both to inhibit drug-seeking during unavailability and promote seeking during availability^41,43^. Thus, the prominent recruitment of IT neurons at DA/DU transitions in early IntA SA may support acquisition of the task structure, while persistent encoding of DA/DU transitions into late IntA SA by IT neurons may contribute to the temporal organization of fentanyl-seeking displayed by our well-trained rats, in which the majority of fentanyl seeking and intake occurs in binge-like bursts primarily within the first 60 seconds of each DA period. Recent work has emphasized the importance of discriminative stimuli in the regulation of addiction-like behaviors; indeed, presentation of a discriminative stimulus can promote drug-seeking even more effectively that a conditioned stimulus^76,77^.

Our data indicate that the responses of prelimbic IT neurons to fentanyl availability and infusions across the IntA SA paradigm correlate with self-administration behaviors. We found that IT activation at DA onset was positively correlated with not only ongoing self-administration behaviors, but also with overall measures of addiction severity, such that the initial responsiveness of IT neurons to fentanyl availability was predictive of subsequent patterns of drug intake. Indeed, low-risk animals in our photometry cohort displayed minimal response to DA onset across IntA SA. We observed that calcium transients associated with lever extension at session initiation and non-contingent cues all increased across SA timepoints, suggesting greater responsiveness by IT neurons to unexpected presentation of fentanyl-associated stimuli over the course of our fentanyl SA paradigm. In contrast, following initial robust drug-associated activation of IT neurons during early training sessions, IT calcium activity negatively correlated with aspects of addiction-like behavior, reflecting a prominent pattern of plasticity in which IT responses to fentanyl infusions and contingent cue presentations diminished in magnitude across days of IntA SA.

Our observation of distinct and dynamic neural encoding of contingent and non-contingent drug-associated cues aligns with a recent longitudinal study of dopamine release within the NAcC, in which non-contingent presentation of a conditioned stimulus elicited phasic dopamine release in NAcC that increased over weeks of cocaine SA, while in parallel, dopamine transients that accompanied self-administered cocaine infusions and contingent cue presentations diminished^78^. Along with the NAcC, the prelimbic PFC receives dopaminergic input important for cue-reward learning and phasic dopamine release within PFC accompanies the presentation of reward-predictive cues^79^. IT neurons express the D1R dopamine receptor ^45,47,80^, and D1R-expressing prelimbic neurons projecting to NAcC exhibit enhanced intrinsic excitability and postsynaptic strength following abstinence from opioid self-administration^81^, consistent with a reported role for prelimbic D1R function in cue-induced reinstatement of opioid seeking^38^. Our observation of diverging IT responsiveness to contingent and non-contingent presentation of fentanyl-associated stimuli may therefore reflect a common pattern of plasticity across multiple nodes of the cortico-basal ganglia network involving dynamic regulation of behavior by the dopaminergic system. Enhanced cortical response to non-contingent cues may support incentive sensitization, whereby drug-associated stimuli drive increasing levels of drug-seeking over time, while decreased response to contingent cues may contribute to evolving experience of satiety and escalation of drug-taking^78^. We observed a prominent IT calcium response evoked by the non-contingent conditioned cue at the onset of cued-reinstatement (CR) sessions that was significantly larger than that observed during lever extension in preceding extinction sessions or in response to prior non-contingent cue presentations. Thus, IT plasticity may enhance the cue-evoked activation of corticostriatal signaling that drives reinstated drug-seeking^40,48,50,52,53,58,59,62,81,82^.

It is intriguing that we observe heterogenous IT event-related transients around fentanyl infusions during well-trained SA sessions, as well as during missed infusions in CR sessions. A subset of rats exhibited a sharp decrease in GCaMP6s fluorescence not present in earlier phases of the behavioral paradigm. While such decreases in fluorescence may reflect synchronous decreases in firing rate in the IT population^83,84^, the significance of such a decrease from a neural coding perspective is unclear. As fiber photometry can only report an aggregate population-level fluorescence, and the normalized amplitude of event-related transients is measured relative to a baseline preceding each event, it is possible that in these animals IT neurons are firing at relatively high rates leading up to lever presses, and are subsequently subject to powerful feedback inhibition once the instrumental action of lever pressing has been performed. Although we note that all type 2 responses were observed in high-risk rats, the relatively small sample size of our fiber photometry cohort precludes meaningful correlation of type 1 and type 2 IT response patterns with specific features of fentanyl self-administration behavior. Future investigations should further interrogate individual variation in the encoding of fentanyl-associated stimuli and relation of such encoding to the severity of addiction-like behaviors.

In summary, our data suggest that robust acquisition of operant contingencies and responsiveness to drug-associated cues is a core feature of the severe high-risk addiction phenotype in our model. Drug-cue associations can guide addiction-like behavior through the corticostriatal pathway in which PFC projection neurons provide crucial glutamatergic input to targets such as the NAcC medium spiny neurons^48,50,52,53,58–62^. While the neurological and genetic underpinnings of individual variation in addiction susceptibility in humans remain largely unknown, alterations to the PFC resulting from chronic drug use have been widely reported^21–25^. Our fentanyl IntA SA paradigm effectively produces a defined subpopulation of rats with a severe addiction-like behavioral phenotype and thus provides an opportunity to interrogate the contribution of specific cortical neuronal populations and circuits to maladaptive opioid-seeking and use.

---

**Supplemental Figure 1.**
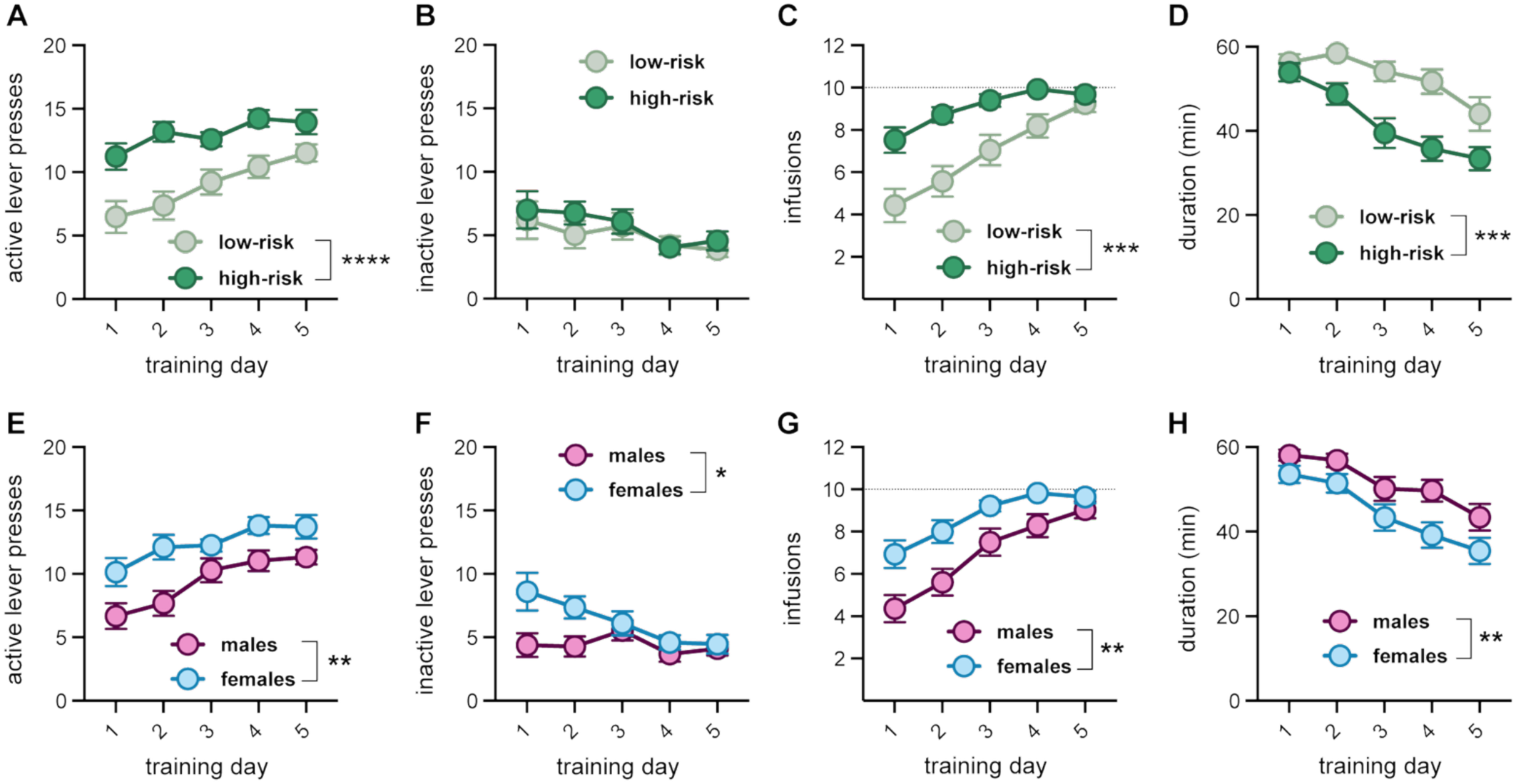
Rats quickly acquire fentanyl self-administration behavior and display lever discrimination and escalating intake. **(A – D).** Animals later classified as low-risk or high-risk differed in their behavior over the training days. **(A, C)** High-risk rats exhibited a greater number of active lever presses and infusions with significant differences during the first four days of training (main effect of severity: F_(1, 44)_ = 22.27, P < 0.0001 for active lever presses; F_(1, 44)_ = 16.67, P = 0.0002 for infusions, two-way RM ANOVA). **(B)** No differences were observed in inactive lever pressing across addiction severity groups**. (D)** High-risk rats completed training sessions faster than low-risk animals (main effect of severity, low- versus high-risk: F_(1, 44)_ = 19.45, P < 0.0001; two-way RM ANOVA). **(E – H)** Male and female rats exhibited differences in self-administration behavior during training days. **(E, G)** Female rats presses the active lever a greater number of times and earned more infusions than males (main effect of sex: F_(1, 54)_ = 12.04, P = 0.0010 for active lever presses; F_(1, 54)_ = 11.20, P = 0.0015 for infusions, two-way RM ANOVA). **(F)** Female rats also showed greater inactive lever pressing (main effect of sex: F_(1, 54)_ = 5.826, P = 0.0192, two-way RM ANOVA). **(H)** Female rats completed training sessions more quickly than males (main effect of sex: F_(1, 54)_ = 9.108, P = 0.0039, two-way RM ANOVA). N = 51 rats total, 21 low-risk, 25 high-risk, 27 male, 24 female.

**Supplemental Figure 2.**
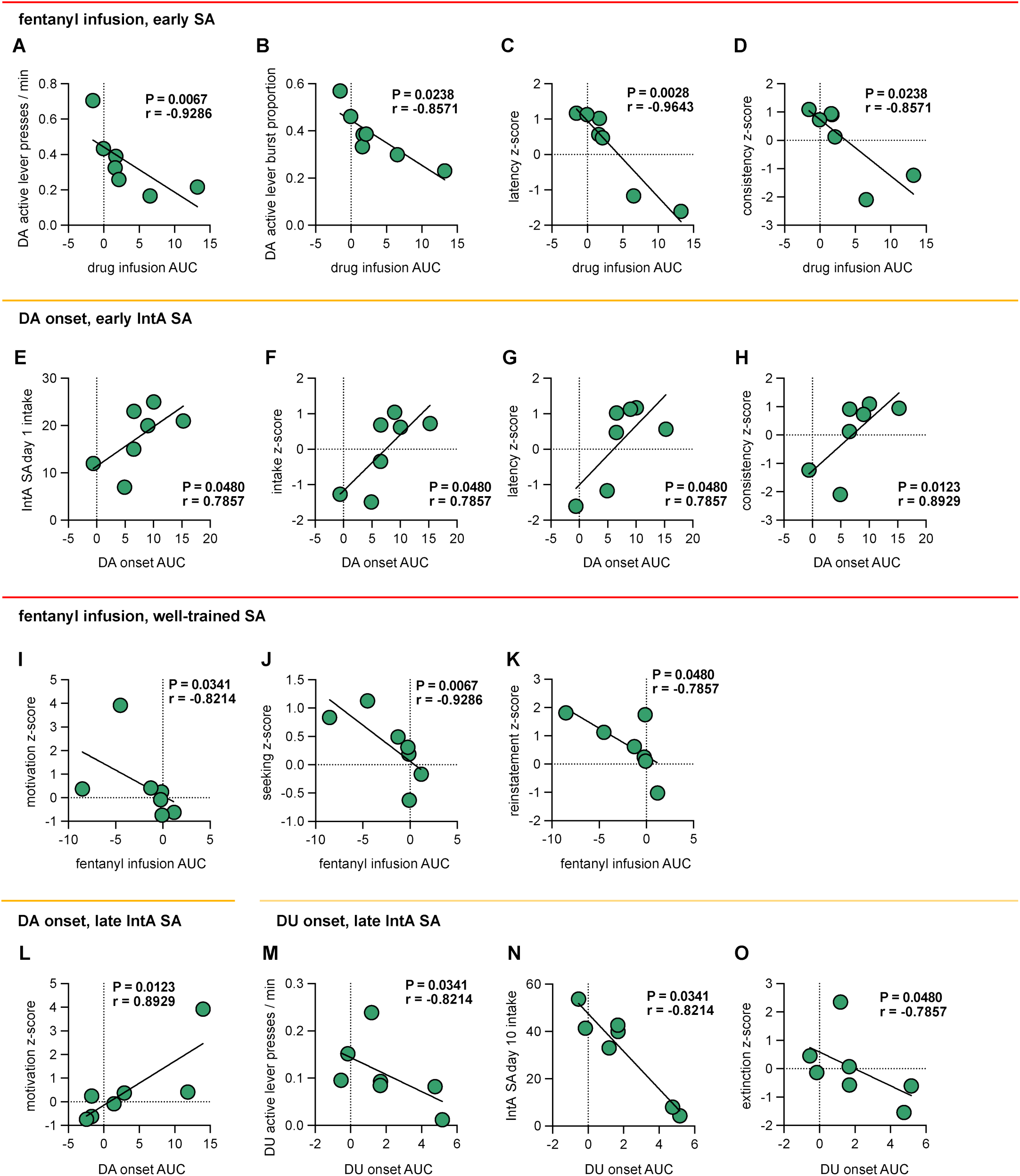
IT event-related calcium transients (ERTs) associated with fentanyl infusions and onset of drug availability correlate significantly with measures of addiction-like behavior. During early SA photometry recording sessions, the magnitude of the fentanyl infusion ERT negatively correlates with **(A)** the rate of active lever pressing, **(B)** the proportion of active lever presses occurring within bursts, **(C)** the latency z-score, **(D)** and the consistency z-score. On the first day of IntA SA, the magnitude of IT ERTs occurring at the onset of drug availability positively correlated with **(E)** IntA SA day 1 fentanyl intake, **(F)** overall intake z-score, **(G)** latency z-score, and **(H)** consistency z-score. In well-trained SA timepoints, fentanyl infusion ERTs correlated negatively with several metrics used for severity classification including **(I)** the motivation z-score, **(J)** seeking z-score, and **(K)** reinstatement z-score. In late IntA SA, DA onset ERTs positively correlated with **(L)** the motivation z-score, while the DU onset ERT negatively correlated with the **(M)** rate of active lever pressing during DU periods, **(N)** the number of fentanyl infusions on day 10 of IntA SA, and **(O)** the extinction z-score.

**Supplemental Figure 3.**
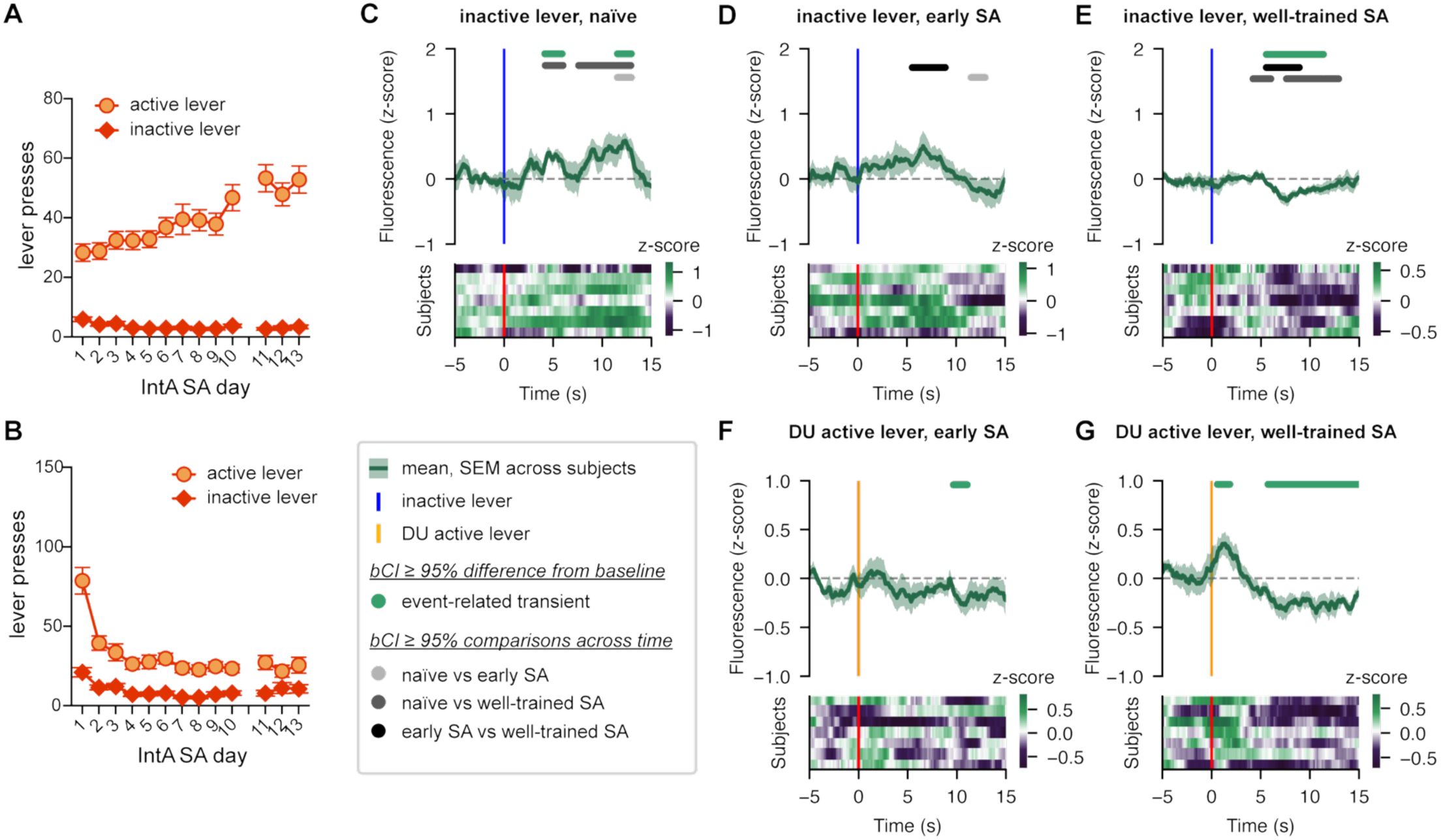
(A) Active lever presses gradually escalated across days of IntA SA while inactive lever pressing remained at minimal levels. **(B)** Rats quickly decrease unrewarded active lever pressing during DU periods across days of IntA SA. **(C – E)** Inactive lever press ERTs evolve across naïve, early SA, and well-trained SA timepoints such that an initial elevation in fluorescence diminishes and a delayed inhibitory response appears in well-trained SA. **(F, G)** DU active lever press ERTs become more apparent from early SA to well-trained SA timepoints.

**Supplemental Figure 4.**
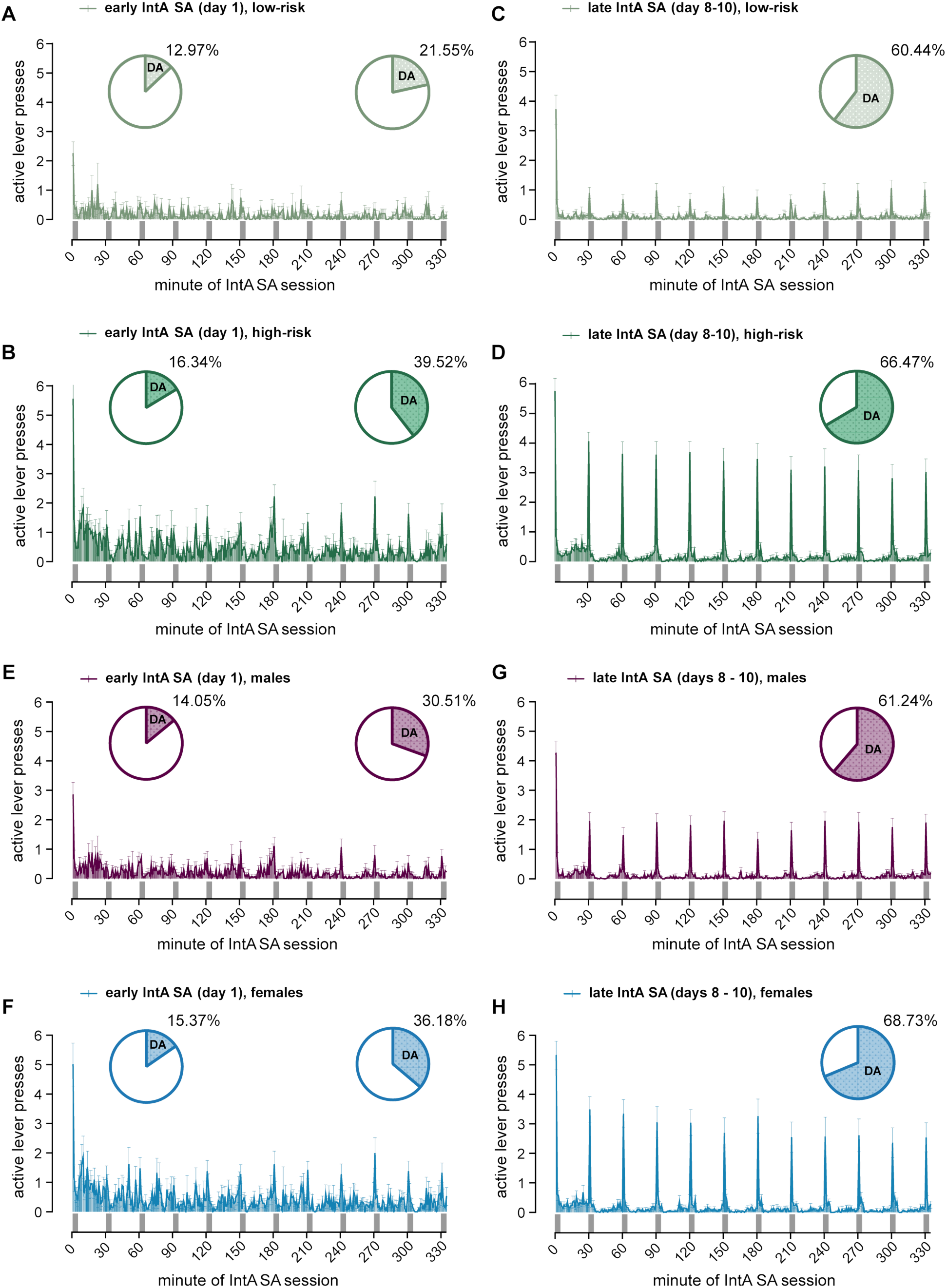
Active lever press patterns in IntA SA drug available and drug unavailable periods across addiction severity and sex. Active lever presses were averaged into one minute bins to illustrate differences in the pattern of drug seeking between low- and high-risk rats, and between male and female rats, both during early IntA SA (day 1) and late IntA SA (days 8-10). **(A, B)** On the first day of IntA SA, high-risk rats exhibited greater active lever pressing than low-risk rats, and near the end of the first day high-risk animals began to concentrate their lever presses within available periods. **(C, D)** During late IntA SA, high-risk rats exhibited greater active lever pressing than low-risk rats **(E, F)** On the first day of IntA SA, female rats exhibited greater active lever pressing than male rats, and near the end of the first day both males and females began to concentrate their lever presses within available periods. **(G, H)** During late IntA SA, female rats exhibited greater active lever pressing than male rats, particularly within available periods.

